# Antibody-free enzyme-assisted chemical labeling for detection of transcriptome-wide *N*^6^-methyladenosine

**DOI:** 10.1101/790683

**Authors:** Ye Wang, Yu Xiao, Shunqing Dong, Qiong Yu, Guifang Jia

## Abstract

The inert chemical property of RNA modification *N*^6^-methyladenosine (m^6^A) makes it very challenging to detect, and all of the transcriptome-wide m^6^A detection methods rely on m^6^A-antibody immunoprecipitation. However, their results are dependent on the quality and specificity of antibodies. Although the endoribonuclease-based single-base m^6^A sequencing is antibody-free, it maps only 16~25% sites. Here, we present an antibody-free, FTO-assisted chemical labeling method termed m^6^A-SEAL for m^6^A detection. We applied m^6^A-SEAL to profile m^6^A landscapes in human and plant, which had good overlaps with antibody-based results and displayed the known m^6^A distribution features in transcriptome. Comparison with all available m^6^A sequencing methods and specific m^6^A sites validation by SELECT, we demonstrated that m^6^A-SEAL has good sensitivity, specificity, and reliability for transcriptome-wide detection of m^6^A. Given its tagging ability and FTO’s oxidation property, m^6^A-SEAL enables many applications like enrichment, imaging, and sequencing techniques to drive future functional studies of m^6^A and other modifications.

## Introduction

*N*^6^-methyladenosine (m^6^A) is the most prevalent chemical modification in mRNA and lncRNA in eukaryotes^1^. This epitranscriptomic mark has essential regulatory functions in RNA processing and metabolism, and so-called writer, eraser, and reader proteins for the mark have been identified. m^6^A is installed by an m^6^A writer complex—the core subunits have been identified as METTL3, METTL14, and WTAP in human^2,3^—and is erased by the AlkB family dioxygenases (e.g., FTO and ALKBH5 in human)^4,5^. m^6^A is read by m^6^A-binding proteins, and such reading has been shown to affect RNA processing in a way that regulates cell physiology^6–11^. Moreover, aberrant m^6^A methylation has been associated with various diseases^12–15^. Therefore, mapping of the transcriptomic distribution of m^6^A can deepen our basic understanding of epitranscriptomic metabolism on cellular physiology and can potentially define novel targets for fighting diseases.

Due to the inert reactivity of the methyl group of m^6^A, transcriptome-wide m^6^A detection methods must rely on m^6^A-antibody immunoprecipitation (m^6^A-IP). m^6^A-seq (also termed MeRIP-seq) is the first developed and presently most widely used method for m^6^A detection^16,17^: it combines m^6^A-IP of fragment mRNAs with high-throughput sequencing to locate m^6^A sites within RNA segments of approximately 200 nucleotides. More recently developed m^6^A detection methods have adapted protocol advantage steps from cross-linking immunoprecipitation RNA target identification methods (e.g., iCLIP and PAR-CLIP) to develop the miCLIP (or m^6^A-CLIP) and PA-m^6^A-seq methods^18–20^, which both can map m^6^A marks at higher resolution than m^6^A-seq based on antibody crosslinking (via either UV or 4-thiouridine). Another alternative m^6^A detection method is called m^6^A-LAIC-seq^21^; this relies on m^6^A-IP of full-length poly(A)^+^ RNA and quantifies methylated transcripts vs. non-methylated transcripts in the immunoprecipitated RNA population. More recently, an antibody-free m^6^A-seq method (m^6^A-REF-seq and MAZTER-seq) using RNA endoribonuclease MazF drew single-base resolution m^6^ACA map on transcriptome; but it can only identify 16~25% of m^6^A sites^22,23^. Notably, most of these methods are dependent on the specificity of anti-m^6^A antibodies, thus emphasizing the strong desirability of developing transcriptome-wide and antibody-free methods to facilitate the routine drawing of m^6^A atlases for many cell and even organismal contexts. However, it is a very challenging to develop a chemical-assisted sequencing method for m^6^A detection due to the inert chemical property of m^6^A.

Here, we present a dithiolsitol (DTT)-mediated thiol-addition chemical reaction that transfers unstable *N*^6^-hydroxymethyladenosine (hm^6^A) to more stable *N*^6^-dithiolsitolmethyladenosine (dm^6^A). Combination with FTO’s enzymatic oxidation of m^6^A on RNA to hm^6^A, we developed FTO-assisted m^6^A selective chemical labeling method (termed as m^6^A-SEAL) for transcriptome-wide detection of m^6^A. m^6^A-SEAL identified 7,785 m^6^A sites in human HEK293T and 12,297 m^6^A sites in rice seedling, which had good overlaps with antibody-based results and displayed the known m^6^A distribution features in transcriptome. Comparison with all available m^6^A sequencing methods, m^6^A-SEAL-unique sites have the same m^6^A distribution profiles along transcripts as the common sites, however, the unique sites from all antibody-based m^6^A-seq methods display distinct features. Eight specific sites (five m^6^A sites and three unmethylated sites) were validated by SELECT method, in line with m^6^A-SEAL results, but not other m^6^A-seq methods. These results reveal that m^6^A-SEAL is a reliable and robust method for transcriptome-wide detection of m^6^A.

## Results

### FTO-assisted selective chemical labeling of m^6^A in RNA

The inert chemical property of m^6^A makes it very challenging to develop a chemical-assisted sequencing method for m^6^A detection. To overcome this challenge, we developed a FTO-assisted m^6^A selective chemical labeling method (m^6^A-SEAL) (Fig. 1). Consider that RNA demethylase FTO oxidizes m^6^A to hm^6^A in 5 min and further slowly oxidizes this hm^6^A to *N*^6^-formyladenosine (f^6^A)^24^. These two intermediates are not stable in aqueous solution (only ~3-hour half-life). Fundamentally, our strategy coupled FTO’s enzymatic oxidation of m^6^A marks on RNA to hm^6^A with a dithiolsitol (DTT)-mediated thiol-addition reaction to install a free sulfhydryl and generate substantially more stable *N*^6^-dithiolsitolmethyladenosine (dm^6^A)-marked RNA species from biological samples (Fig. 1a). Having overcome the limited reactivity of m^6^A marked RNA species, we could exploit the free sulfhydryl group present on dm^6^A to install a variety of tags (e.g., biotin and fluorophores) through reaction with methanethiosulfonate (MTSEA), thereby easily facilitating applications including detection, sequencing and imaging (Fig. 1b).

**Fig. 1.**
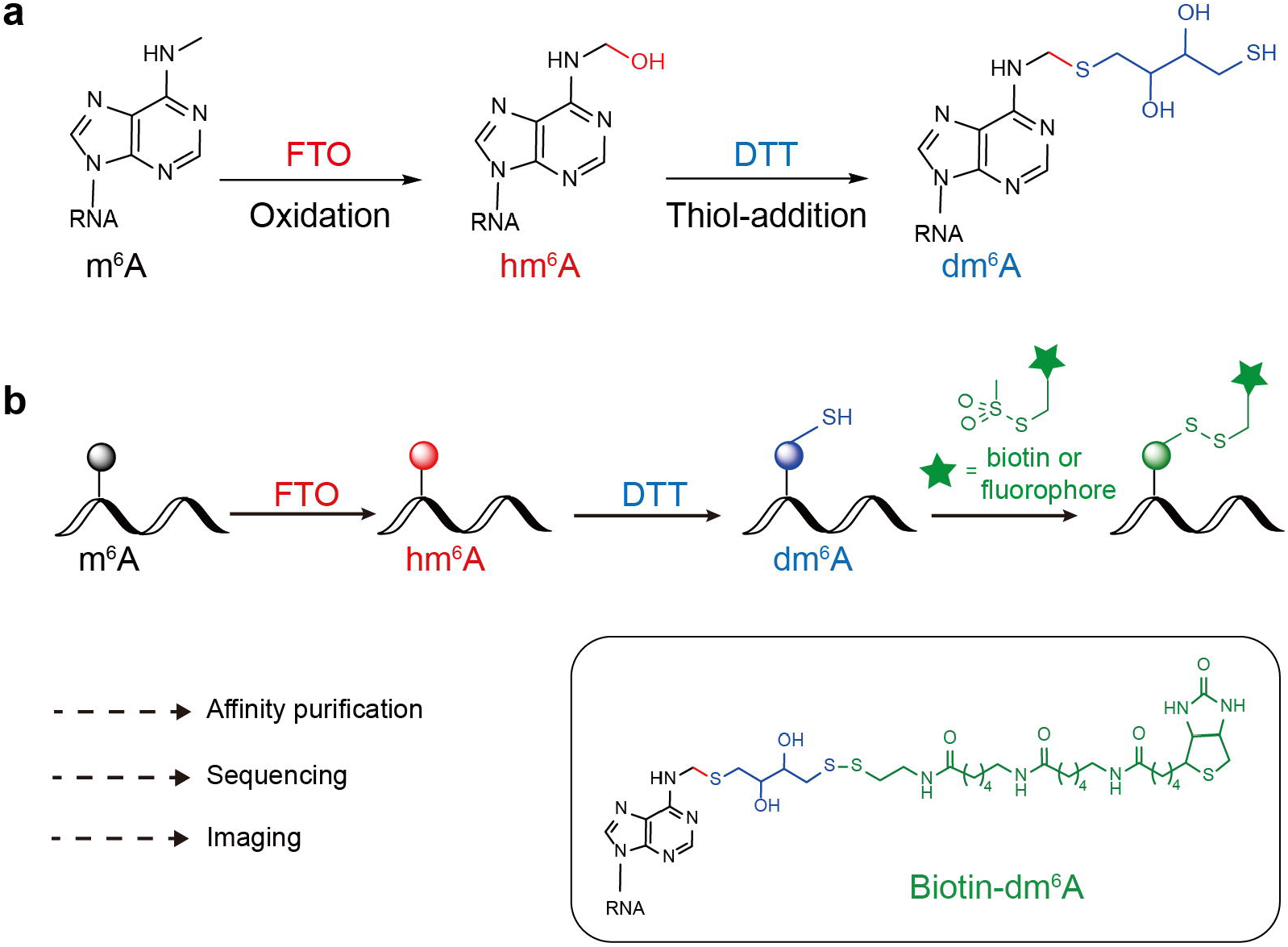
FTO-assisted selective chemical labeling of m^6^A. **a**, FTO oxidation of m^6^A to hm^6^A and a DTT-mediated thiol-addition reaction to convert unstable hm^6^A to the more stable dm^6^A. **b**, Schematic diagram of m^6^A-SEAL. m^6^A in RNA can be sequentially converted to hm^6^A and dm^6^A. The free sulfhydryl group on dm^6^A reacts with MTSEA to install a variety of tags (e.g., biotin and fluorophore), thereby easily facilitating applications including affinity purification, sequencing, and imaging.

We initially explored a variety of chemistries for the modification of hm^6^A using a model system based on chemically synthesized hm^6^A single nucleoside and synthetic m^6^A-modified RNA oligonucleotide. Inspired by the aminomethylation product in formaldehyde induced DNA-protein crosslinking^25^, we quickly narrowed our attention to the use of sulfhydryl reagents to convert the unstable hemiaminal structure of hm^6^A into a more stable thiol-aminal structure, and found that DTT performed very well for derivatizing hm^6^A nucleoside in aqueous solution (Supplementary Fig. 1 and Supplementary Note 1). Other sulfhydryl reagents were also able to react with hm^6^A single nucleoside (Supplementary Fig. 2). Given the additional attractive benefit of a second free sulfhydryl provided by DTT, which should facilitate subsequent functional tagging for downstream applications, we chose DTT for further study. We used mass spectrometry and NMR to confirm the chemical structures of dm^6^A produced by our DTT treatment of hm^6^A nucleoside (Supplementary Note 2). Presumably, the thiol-addition reaction on hm^6^A is accomplished via [1, 2] addition of imine, but detailed mechanism remains to be fully elucidated (Supplementary Fig. 3).

Optimization showed that the yield of the DTT-mediated thiol-addition on hm^6^A nucleoside reached as high as 80% under mild acidic condition (~pH 4.0) at 37 °C over 3 h (Supplementary Figs. 1 and 4). We evaluated the stability of the dm^6^A nucleoside at various pH conditions. Our finding that dm^6^A had a much longer half-life (182 h) in neutral aqueous solution than hm^6^A (3 h, as previously reported^24^) (Supplementary Fig. 5) supported the basic stability-enhancing concept of our method’s design and suggested that the two-reaction process was likely practicable with biological samples.

Having optimized the conversion of dm^6^A from hm^6^A single nucleoside, we next conducted experiments with synthesized 9mer RNA oligonucleotide (model RNA) bearing m^6^A modification at the 5’ end. Initial testing with the model RNA identified the optimal FTO oxidation and DTT-mediated thiol-addition reaction conditions to obtain the highest yield of the conversation of dm^6^A from m^6^A. Importantly, our choice to position the m^6^A mark at the 5’ end in the model RNA ultimately facilitated precise HPLC-based quantification of the products from two reactions: the FTO-catalyzed oxidation of m^6^A to hm^6^A and the DTT-mediated thiol-addition of hm^6^A to dm^6^A. Specifically, the nucleoside at the 5’ end of model RNA can be selectively released via nuclease P1 treatment in neutral pH condition within 15 min, which efficiently reduced the degradation of the hm^6^A intermediate during routine RNA digestion procedures. In the optimized two-step reactions, 0.5 nmol of FTO was used to treat with 1 nmol of model RNA for 5 min at pH 7 and 37 °C, generating 60% of hm^6^A formation converted from m^6^A (Supplementary Fig. 6), a yield similar to a previously reported study of FTO kinetics^23^. After ethanol precipitation purification, the purified RNA was subsequently treated with 200 mM of DTT for 3 h at pH 4 and 37 °C, generating 83% of dm^6^A formation from hm^6^A (Fig. 2a and Supplementary Fig. 7).

**Fig. 2.**
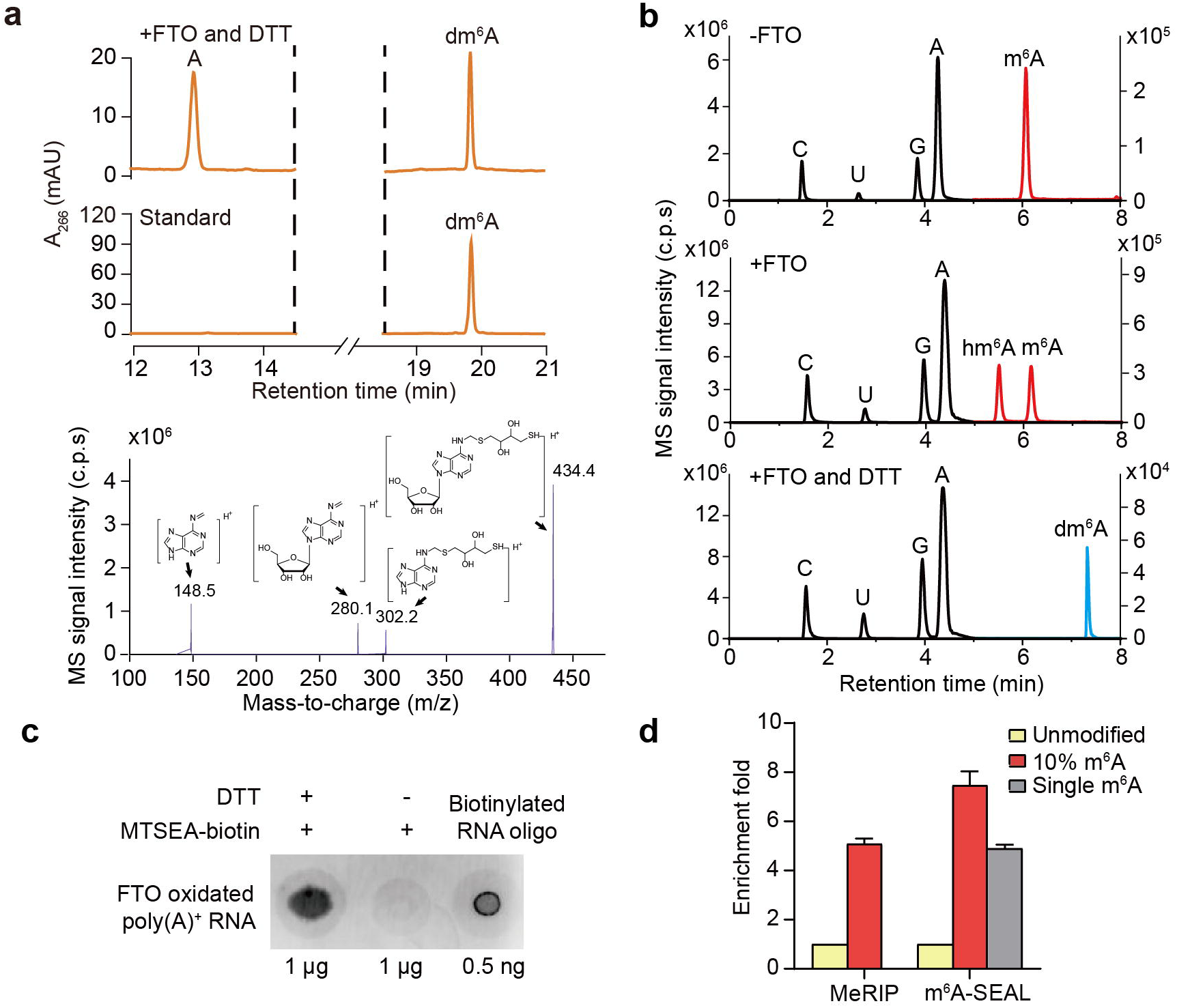
m^6^A-SEAL on model RNA and human poly(A)^+^ RNA. **a**, Upper Panel: HPLC chromatograms showing the formation of dm^6^A on model RNA after FTO oxidation-assisted and DTT-mediated thiol-addition reactions, and the dm^6^A standard. 1 nmol of m^6^A-modified model RNA was treated with 0.5 nmol of FTO for 5 min at pH 7 and 37 °C, followed by incubation with 200 mM DTT for 3 h at pH 4 and 37 °C. Lower Panel: MS/MS profile of dm^6^A nucleoside. **b**, LC-MS/MS chromatograms showing the formation of hm^6^A and dm^6^A on HEK293T poly(A)^+^ RNA after FTO oxidation and FTO oxidation-assisted and DTT-mediated thiol-addition reactions, respectively. 300 ng of HEK293T poly(A)^+^ RNA was treated with 0.2 μM of FTO for 5 min at 37 °C in 100 μl of reaction mixture, followed by incubation with 200 mM DTT for 3 h at pH 4 and 37 °C. The chromatograms of A, U, C, and G are scaled to the left y axis, and the chromatograms of m^6^A, hm^6^A, and dm^6^A are scaled to the right y axis. **c**, Dot-blot assay confirming biotin labeling of m^6^A on HEK293T poly(A)^+^ RNA. A 3’ biotinylated 15mer RNA oligo was used as a positive signal control. **d**, Enrichment of spike-in RNA sequences detected by m^6^A-SEAL-seq and MeRIP-seq. Values represent fold enrichment of IP over input (n=2), normalized to non-m^6^A-modified spike-in. 10% m^6^A, reverse transcribed RNA with 10% of m^6^ATP; single m^6^A, synthetic RNA with a single m^6^A site.

### m^6^A-SEAL sensitively and specifically enriches m^6^A-modified RNA

After validation of m^6^A-SEAL on model RNA, we next tested m^6^A-SEAL on human RNA to optimize the reaction conditions and examine its sensitivity and specificity. First, purified poly(A)^+^ RNA from HEK293T cells was chemically fragmented into 100 nt molecules at 95 °C, which were then treated with FTO (at a variety of concentrations to facilitate optimization). We found that a five-minute incubation at 37 °C with 0.2 μM of FTO resulted in the highest yield (~50%) of hm^6^A using 300 ng of poly(A)^+^ RNA in 100 μL of demethylation reaction, these hm^6^A can be further converted to dm^6^A (yield 69%) under the treatment of DTT (Fig. 2b and Supplementary Fig. 8). Importantly, we detected no noticeable degradation for the fragmented poly(A)^+^ RNA upon this mild acidic condition (Supplementary Fig. 9), emphasizing that the protocol would not substantially affect the quality of RNA and sequencing libraries. The successfully DTT labeled poly(A)^+^ RNA was subsequently reacted with MTSEA-biotin, a commercial thiol-reactive biotin reagent. The labeling of biotin was confirmed with a dot-blot assay (Fig. 2c). No free dm^6^A was detected in LC-MS/MS analysis of the RNA sample following the MTSEA-biotin labeling step indicated very high biotin labeling efficiency (Supplementary Fig. 10).

To test the sensitivity and specificity of m^6^A-SEAL for sequencing applications, we next conducted several spike-in RNAs (Supplementary Note. 3) in poly(A) ^+^ RNA pool to perform m^6^A-SEAL sequencing (m^6^A-SEAL-seq). The high-throughput sequencing results showed that our method facilitated strong enrichment for the spike-in m^6^A-modified sequences (for both singly and multiply m^6^A modified species), which is comparable to MeRIP (Fig. 2d). Importantly, we detected no cross-reactivity for unmodified sequence (Fig. 2d). Therefore, these results support impressive sensitivity and specificity for m^6^A-SEAL based evaluation of m^6^A distribution in biological samples. These spike-in experiments also showed that a five-minute reaction time and 0.2 μM FTO were ideal to obtain the highest enrichment efficiencies of the spike-in m^6^A-modified sequences (Supplementary Fig. 11).

### m^6^A-SEAL identifies transcriptome-wide m^6^A sites in human cells

We subsequently performed m^6^A-SEAL-seq for poly(A)^+^ RNA from HEK293T cells. We detected a total of 14,274 m^6^A peaks within human genes in two biological replicates, and classified the overlapping 7,785 peaks as high confidence sites for subsequent analysis (Supplementary Fig. 12). Several known m^6^A sites in transcripts were consistently identified in two biological replicates of m^6^A-SEAL (Fig. 3a). Note that sequencing on different platforms (and with a variety of analysis modes) yielded satisfactory data in each case (Supplementary Table. 1). The m^6^A sites identified by m^6^A-SEAL-seq revealed strong enrichment for the well-known canonical m^6^A motif RRACH (R=purine, H=A/C/U) (Fig. 3b). The center of m^6^A peaks were highly enriched at the consensus sequence compared to the negative control peaks (Fig. 3c), supporting the specificity of m^6^A-SEAL. The distribution of m^6^A within transcriptomes revealed strong enrichment in the vicinity of stop codons (Fig. 3d), which echoed a typical m^6^A distribution pattern mapped using MeRIP-seq^16,17^. Mapping m^6^A sites to exons revealed that ~89% of m^6^A-modified exons were longer than 400 nt compared to the negative control peaks (p<2.2 × 10^−16^, Wilcoxon test; Fig. 3e), in line with previous finding that m^6^A preferentially locates within long exons^16,20^.

**Fig. 3.**
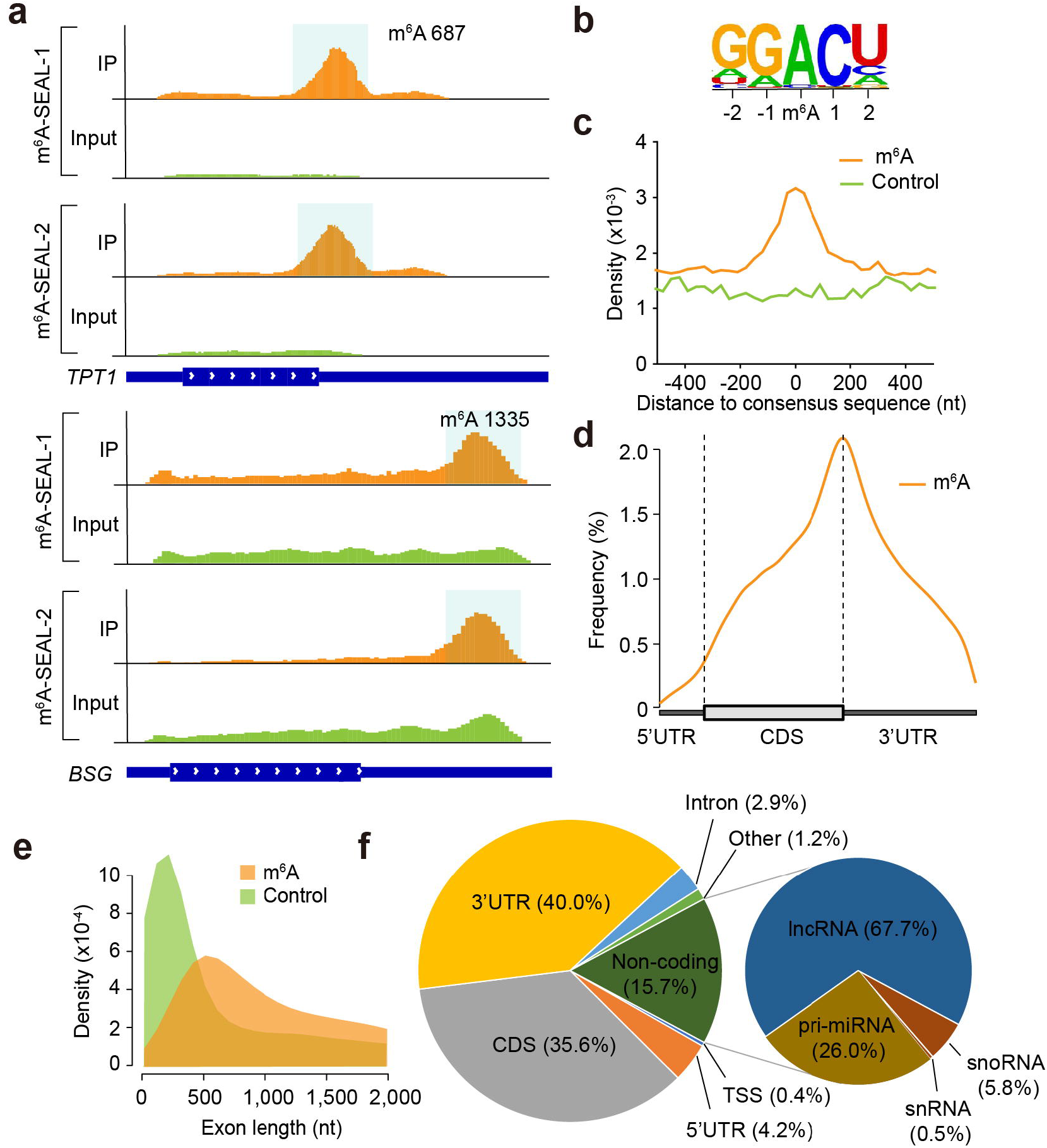
m^6^A-SEAL-seq uncovers the transcriptome-wide m^6^A methylome in HEK293T cells. **a**, Representative views of m^6^A-SEAL-seq identified typical m^6^A peaks on mRNA (*TPT1* and *BSG*). Peaks are highlighted in gray, which contain SCARLET identified m^6^A sites^26^. Transcript architecture is shown below with thin and thick parts respectively representing UTR and CDS. **b**, Canonical RRACH motif identified in m^6^A-SEAL-seq. **c**, Density plots showing the distance of the centre of m^6^A peaks/control peaks between the nearest consensus sequence RRACH. **d**, Metagene profile illustrating the distribution of m^6^A sites identified by m^6^A-SEAL across mRNA segments. **e**, Density plots showing the distribution of m^6^A/control peaks according to their located exon length. **f**, Pie chart presenting the fraction of m^6^A sites identified by m^6^A-SEAL-seq in each non-overlapping RNA segment. **g**, Repeats families (y-axis) ranked by the number of elements containing intergenic m^6^A sites (x-axis).

Closer examination of the distribution of the identified m^6^A sites within regions of transcriptome revealed that the majority of m^6^A (83.1% of 7,785 confident m^6^A sites) were present on coding transcripts with a preference within the coding sequences (CDS) and 3’ untranslated region (3’UTR) (Fig. 3f). 33 m^6^A sites near transcription start sites (TSS, < 150 nt) were regarded as potential *N*^6^,2’-*O*-dimethyladenosine modifications at the cap +1 position (termed cap m^6^A_m_; Fig. 3f), suggesting that our method could capture m^6^A and cap m^6^A_m_ sites simultaneously. 1,222 m^6^A sites (15.7%) belongs to non-coding RNAs, including lncRNA, pri-miRNA, snoRNA, and snRNA (Fig. 3f).

### m^6^A-SEAL confirms that UGUAHH is a plant-specific m^6^A motifs

Having established proof-of-concept for global m^6^A profiling from biological samples, we were next eager to deploy m^6^A-SEAL-seq to explore unsolved scientific questions in the exciting basic research area of epitranscriptomics cellular regulation. MeRIP-seq cannot identify the canonical m^6^A motif RRACH in *Oryza sativa* (rice); instead, a different motif sequence was found but not ascribed with high confidence because of the potential for non-specific antibody binding and the absence of experimental confirmation^27,28^. Considering that our m^6^A-SEAL is a chemically covalent cross-linking method for mapping m^6^A sites, which eliminates the non-specific binding, we addressed this unsettled scientific question by performing m^6^A-SEAL-seq in poly(A)^+^ RNA samples isolated from 60-day-old rice leaves.

12,297 high confidence m^6^A peaks were identified in two biological replicates (Supplementary Fig. 13a). Metagene profiling and the distribution within the non-overlapped segments of transcripts of the m^6^A sites identified by m^6^A-SEAL showed that m^6^A predominantly occurs within 3’UTR (Supplementary Fig. 13b,c), in consistent with the previous MeRIP-seq result in rice^27^. To identify m^6^A motifs, we clustered all confident m^6^A peaks in HOMER (Hypergeometric Optimization of Motif Enrichment) software to search motifs. We did not find the canonical m^6^A motif RRACH in rice; instead, we identified a non-canonical motif UGUAHH (H=A/C/U) with highest significance (Supplementary Fig. 13d). Note that similar motifs were previously found in rice panicles (UGWAMH, W=U or A; M=C or A; H=U, A, or C)^27^ and Arabidopsis (URUAY, R=G or A; Y=U or A)^29^. To further confirm it, we calculated the enrichment of m^6^A-SEAL identified m^6^A peaks to the centre of UGUAHH and RRACH motifs, which showed that m^6^A peaks were strongly enriched at the centre of the UGUAHH motif but not the RRACH motif (Supplementary Fig. 13d).

### m^6^A-SEAL determines m^6^A sites more reliably compared to all available m^6^A sequencing methods

To further evaluate the reliability and performance of m^6^A-SEAL, we compared it with all available m^6^A sequencing methods. We firstly compared our m^6^A-SEAL results identified in HEK293T and rice with MeRIP-seq results. To do this, we used the published MeRIP sequencing data for HEK293T (GSE29714)^2^ and our own constructed MeRIP sequencing data in 60-day-old rice leaves, ~70% of the m^6^A peaks in HEK293T and ~59% rice peaks from m^6^A-SEAL-seq positionally overlapped with peaks from MeRIP-seq (Fig. 4a). We categorized all m^6^A sites from m^6^A-SEAL and MeRIP into three groups for further analysis: common sites (overlapped m^6^A peaks between m^6^A-SEAL and MeRIP), m^6^A-SEAL-unique sites (not identified in MeRIP), and MeRIP-unique sites (not identified in m^6^A-SEAL). Motif search analysis by HOMER revealed that in HEK293T m^6^A-SEAL-unique sites and MeRIP-unique sites had the same m^6^A motif identified in common sites; however, in rice m^6^A-SEAL-unique sites contained the same motif as common sites while MeRIP-unique sites can not identify any significant motif (Fig. 4b). Metagene profilings showed that m^6^A-SEAL-unique sites both in HEK293T and rice had the same m^6^A distribution feature as common sites, however, MeRIP-unique sites presented a distinct feature (Fig. 4c). We next compared our m^6^A-SEAL results with the published miCLIP results in HEK293T, an antibody and CLIP-based single-base m^6^A sequencing method^18^. m^6^A sites identified by either cross-linking-induced truncation sites in miCLIP (miCLIP-CITS) or cross-linking-induced mutation sites in miCLIP (miCLIP-CIMS) had 40~50% of overlap with that of m^6^A-SEAL (Supplementary Fig. 14). Further metagene profilings revealed that both miCLIP-CITS-unique sites and miCLIP-CIMS-unique sites exhibited distinct m^6^A distribution features from the common m^6^A sites (Fig. 4c). These results suggest antibody-based m^6^A sequencing methods might have non-specific antibody binding effect.

**Fig. 4.**
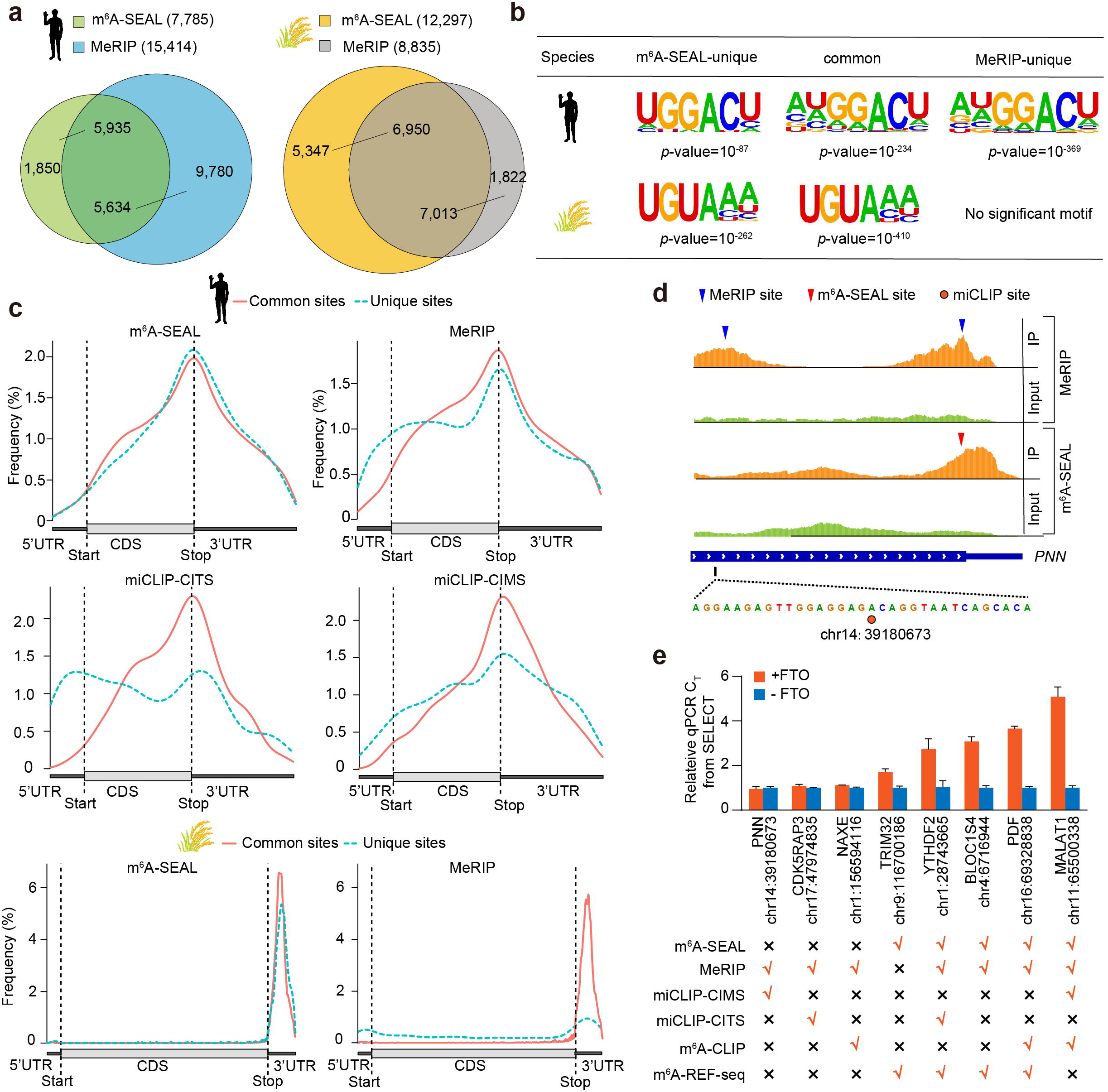
Comparison of m^6^A methylone in human HEK293T cells and rice detected by m^6^A-SEAL-seq and all available m^6^A-seq methods. **a**, Overlap of m^6^A peaks identified by m^6^A-SEAL-seq and MeRIP-seq. **b**, Human and rice m^6^A motif enriched in common m^6^A sites (identified in both m^6^A-SEAL and MeRIP-seq), m^6^A-SEAL-unique sites (not present in MeRIP), and MeRIP-unique sites (not present in m^6^A-SEAL). **c**, Metagene profile of the distribution of common m^6^A sites (identified in both m^6^A-SEAL and MeRIP-seq (or miCLIP)) and respective unique m^6^A sites across human and rice mRNA segments. **d**, Representative view of m^6^A sites on mRNA *PNN* identified by m^6^A-SEAL, MeRIP, and miCLIP. **e**, SELECT validation of eight individual sites detected by different m^6^A-seq methods in HEK293T. Eight sites are validated as three unmethylated sites and five m^6^A sites by SELECT. Bar plots showing the relative threshold cycle (C_T_) of qPCR for detecting the m^6^A site (normalized by input control site). Data are presented as means ±SD, n=2 biological replicates ×3 technical replicates. The m^6^A sites of MeRIP, miCLIP-CIMS, miCLIP-CITS, m^6^A-CLIP, and m^6^A-REF-seq are obtained from the literatures.

To further demonstrate the reliability of m^6^A-SEAL, we picked eight sites—five m^6^A sites and three unmethylated sites—from m^6^A-SEAL results for validation using SELECT method we developed previously (Fig. 4d and Supplementary Fig. 15). SELECT is an elongation and ligation-based qPCR amplification method for detection of m^6^A position and fraction in mRNA/lncRNA at single-nucleoside resolution^30^. SELECT method confirmed these five m^6^A sites and three unmethylated sites, which were consistent with m^6^A-SEAL results (Fig. 4e). SELECT validation results showed that all antibody-based m^6^A sequencing methods had high false positive rates. In total eight sites, MeRIP had three false positives and one false negatives, (Fig. 4d,e and Supplementary Fig. 15). The antibody-based single-base m^6^A sequencing methods (miCLIP and m^6^A-CLIP) displayed low ability for mapping m^6^A sites and high false positive rates. In total eight sites, miCLIP-CIMS and miCLIP-CITS together identified four m^6^A sites, but only half of them were positives; m^6^A-CLIP identified three m^6^A sites, but one of them was false positives (Fig. 4d,e and Supplementary Fig. 15). m^6^A-REF-seq as an antibody-free and endoribonuclease-based single-base m^6^A detection method exhibited good reliability on ACA motif, however, it was powerless to map other m^6^A-modified motifs (for instance, m^6^A-REF-seq can not identify m^6^A-modified ACU motif in *MALAT1* 2577 site (chr11: 65500338)) (Fig. 4e and Supplementary Fig. 15). Collectively, m^6^A-SEAL as a chemical labelling method has higher reliability for detection of m^6^A sites.

## Discussion

The epitranscriptomic modification m^6^A regulates numerous biological processes and diseases. It is desirable to develop a chemical-assisted m^6^A sequencing to reduce the non-specific antibody binding effect. In this work, we report an FTO-assisted m^6^A selective chemical labeling method (m^6^A-SEAL) for transcriptome-wide detection of m^6^A. This method combines two reactions in mild conditions: FTO-assisted oxidation of m^6^A to yield an unstable intermediate hm^6^A and DTT-mediated thiol-addition reaction to stabilize and enable functionalization of this intermediate. m^6^A-SEAL has multiple advantages: it is an antibody-free covalent cross-linking method, requires only commercial available reagents (Note that FTO protein is commercial available or can be easily purified in laboratory; DTT is a common biochemical reagent.), and used available bioinformatics tools for data processing. We demonstrated that this cost-effective, antibody-free method has good sensitivity, specificity, and reliability for transcriptome-wide detection of m^6^A.

Transcriptome-wide m^6^A sites in both human HEK293T and rice identified by m^6^A-SEAL had good overlaps with MeRIP results and displayed the known m^6^A distribution features in transcriptome. Compared with all available m^6^A sequencing methods, m^6^A-SEAL-unique sites performed the same m^6^A distribution feature as common sites, however, MeRIP-unique sites and miCLIP-unique sites showed distinct m^6^A distribution profiles. Single-base m^6^A sites validation by SELECT showed that all antibody-based m^6^A sequencing methods read unmethylated sites as m^6^A sites, further confirming the non-specific antibody binding leads to false positives. The ΔC_T_ values of SELECT between FTO treatment (+FTO) and without FTO treatment (-FTO) indicate the m^6^A methylation level^30^. SELECT results showed that the methylation fraction of m^6^A 2605 site on *TRIM32* transcript (chr9: 116700186) is the lowest among these five identified m^6^A sites. All antibody-based m^6^A sequencing methods failed to map this site, but m^6^A-SEAL succeed, indicating m^6^A-SEAL has higher sensitivity. m^6^A-REF-seq is an FTO-assisted endoribonuclease MazF-dependent single-base m^6^A sequencing method that only reads m^6^A sites in ACA motif^22^. Note that the three unmethylated sites and one *MALAT1* m^6^A site belong to non-ACA motif. m^6^A-REF-seq is powerless to detect these sites, however, m^6^A-SEAL can distinguish whether these sites are modified or not.

Cap m^6^A_m_ is a new identified reversible RNA modification in human and mouse poly(A)^+^ RNA^31–34^. m^6^A-SEAL identified 33 m^6^A sites near TSS, which could be potential cap m^6^A_m_ sites. As known that FTO can oxidize cap m^6^A_m_ to hm^6^A_m_ and FTO demethylates cap m^6^A_m_ *in vitro* with a higher efficiency over that of internal m^6^A^31^, by optimization of the FTO oxidation conditions, the procedure of m^6^A-SEAL could be used for detection of cap m^6^A_m_ modification. Additionally, FTO also oxidizes DNA *N*^6^-methyldeoxyadenosine (6mA) to *N*^6^-hydroxymethyldeoxyadenosine (6hmA)^4,24^. DNA 6mA is a new emerging epigenetic modification with low abundance^35–39^. m^6^A-SEAL could be optimized to be a potential sensitive and reliable method for detection of DNA 6mA modification.

hm^6^A is an unstable modification that is present in poly(A)^+^ RNA isolated from human cells and mouse tissues^24^. Its ~3-hour half-life is comparable to the average half-life of mRNA in mammalian cells^40^; whether hm^6^A plays a biological function in cells is unknown because it is hard to be detected and studied. Without FTO oxidation step, DTT-mediated hm^6^A labeling method could be adapted to detect this unstable modification in transcriptome-wide manner. Furthermore, m^6^A-SEAL converts the inert m^6^A to dm^6^A with a free sulfhydryl, which can be installed a variety of tags (e.g., biotin and fluorophores) through reaction with MTSEA. Here we have demonstrated one application that we installed biotin tag on m^6^A for enrichment and sequencing. We anticipate that m^6^A-SEAL method could be modified to image cellular m^6^A modification by installation of fluorophores on cellular m^6^A.

In summary, we developed m^6^A-SEAL method, which couples FTO’s enzymatic oxidation of m^6^A to the unstable intermediate hm^6^A with DTT-mediated thiol-addition reaction to generate a more stable dm^6^A with a sulfhydryl group that facilitates simple installation of functional application tags like biotin. We demonstrated that m^6^A-SEAL is a robust and reliable method for detection and functional studies of m^6^A. Considering FTO’s oxidation property on multiple modifications and the tagging ability of dm^6^A, m^6^A-SEAL with optimization could be adapted to detect other modifications and image cellular m^6^A modification.

## Acknowledgements

We would like to acknowledge S. Liu, J. Meng and H. Liu for helping with the data analysis. This work was supported by the National Basic Research Program of China (2017YFA0505201) and the National Natural Science Foundation of China (nos. 21822702, 21432002, and 21820102008).

## Author contributions

G.J. and Y.W. conceived the project and designed the experiments; Y.W. performed the experiments and data analysis with the help of S.D., Q.Y., and Y. X.; G.J. and Y.W. wrote the manuscript.

## Additional information

**Correspondence and requests for materials** should be addressed to G.J.

## Methods

### Synthesis of model RNA oligonucleotides

9mer model RNA containing m^6^A (5’-m^6^ACUGACUAG-3’) was synthesized on ABI Expedite 8909 nucleic acid synthesis system using commercially standard RNA phosphoramidites and m^6^A phosphoramidite (Glen Research) in DMT-ON model. The phosphate backbone of the first three bases on 5-terminal and the last three bases on 3-terminal were thiolated by Beaucage reagent (Glen Research) in order to prevent RNase degradation. Purification of oligo nucleotide is carried out by Glen-Pak cartridges (Glen Research).

### Expression and purification of recombinant human FTO protein

Protein expression was performed as previously described with minor modifications^41^. Briefly, human N-truncated human FTO protein (NP_001073901.1, 31-505aa) was subcloned into pET28a vector. Plasmid was transformed into *Escherichia coli* BL21(DE3) and grew on LB agar plate with 50 mg/L kanamycin. After being inoculated to 1 L LB medium that contained 50 mg/L kanamycin, bacteria were cultured under 37°C, 220 r.p.m in shaker until OD_600_ reached 1.0. The expression of protein were induced by 0.5 mM IPTG at 16°C for overnight. Cell pellets were harvested and resuspended in 20 mL Buffer A (10 mM Tris pH 7.9, 150 mM NaCl) and sonicated on ice. Lysate was centrifugated for 30 min at 13000g and 4°C. Supernatant was collected and filtered by 0.22 μm filter (Millipore) then loaded on Ni-NTA column (GE Healthcare). After washed by 20 mL Buffer A and then 20 mL 8% Buffer B, protein was eluted by Buffer B (10 mM Tris pH 7.9, 150 mM NaCl, 500 mM imidazole). The collected fraction was then purified by MonoQ ion exchange column(GE Healthcare, Buffer A: 10 mM Tris pH 7.9, 3 mM DTT. Buffer B: 10 mM Tris pH 7.9, 1M NaCl) to remove non-specific nucleic acids binding (0-100% Buffer B gradient) and subsequently purified by Superdex 75 gel-filtration column (GE Healthcare, 10 mM Tris pH 7.9, 150 mM NaCl and 3 mM DTT). Protein was concentrated into 10 mg/mL and 20% glycol was added. Aliquots of protein were frozen by liquid N_2_ then stored in −80°C.

### Human Cell culture and isolation of poly(A)^+^ RNA

HEK 293T/17 cell (CRL-11268, ATCC) was cultured in DMEM medium (Corning) that contains 10% FBS (Cellmax) and 1% Penicillin Streptomycin (Corning) under the condition of 37 °C and 5% CO_2_. Cells were washed once by PBS and harvested for the isolation of total RNA using TRIzol reagent (Magen). Poly(A)^+^ RNA was isolated from total RNA using oligo(dT)_25_ Dynabeads (Thermo Fisher Scientific). RNA concentration was measured by Nanodrop UV-Vis Spectrophotometer (Thermo).

### Rice sample preparation and isolation of poly(A)^+^ RNA

Rice (Nipponbare) seeds were soaked in water in the dark at 30 °C for two days. After that, seedlings were grown for 28 days at 28°C in a soil seed bed. The humidity was 70% with 12h light/12h dark. Then the seedlings were transplanted into pots and grown for 60 days in a greenhouse under normal condition. Rice leaves were harvested and frozen by liquid N_2_, then disrupted using the TissueLyser II (QIAGEN). Total RNA was extracted by TRIzol reagent (Magen) from homogenized samples. Poly(A)^+^ RNA was isolated from total RNA using oligo(dT)_25_ Dynabeads (Thermo Fisher Scientific).

### FTO oxidation of m^6^A to hm^6^A in model RNA

Oxidation of 9-mer model RNA (5 μM) was performed in 300 μM of (NH_4_)_2_Fe(SO_4_)_2_ · 6H_2_O, 2 mM of L-ascorbic acid, 300 μM of α-KG, 100 mM pH=7.0 HEPES, 2.5 μM FTO at 37 °C. After 5 min treatment, the reaction was quenched by 5 mM EDTA and then frozen in liquid N_2_ immediately for HPLC analysis.

### FTO oxidation of m^6^A to hm^6^A in poly(A)^+^ RNA

Poly(A)^+^ RNA was fragmented by Magnesium RNA fragmentation module (NEB). In poly(A) ^+^ RNA oxidation assay, the reaction was performed in 300 μL aliquots of aqueous solution containing 300 μM of (NH_4_)_2_Fe(SO_4_)_2_ · 6H_2_O, 2 mM of L-ascorbic acid, 300 μM of α-KG, 100 mM pH7.0 HEPES, 0.2 μM FTO and 1μg Poly(A) RNA. After the FTO treatment at 37 °C for 5 min, RNA was purified by RNA Clean & Concentrator-5 column (Zymo Research). A large volume of FTO oxidation reaction can be concentrated by lyophilization before column purification.

### DTT labeling of hm^6^A-modified RNA produced by FTO oxidation

hm^6^A-modified RNA converted from m^6^A by FTO oxidation was treated by 200 mM freshly prepared DTT at 37 °C for 3 h in acidic aqueous solution (100 mM HEPES, pH 4.0). The product RNA was purified by ethanol precipitation.

### Biotinylation of DTT-labeled RNA

After ethanol precipitation, DTT treated RNA was washed by 75% ethanol and dissolved in 200 μL biotinylation buffer that contains 100 μM of MTSEA-XX-biotin (Biotum), 100 mM HEPES (pH 7.0), 1 mM EDTA and 20% DMF. The reaction was performed at 25 °C and 800 r.p.m in ThermoMixer for 1h. The product RNA was purified by phenol-chloroform extraction or by RNA Clean & Concentrator-5 column purification.

### Determination of the formation efficiencies of hm^6^A and dm^6^A in model RNA using HPLC analysis

After FTO oxidation and DTT labeling, 5’ nucleoside in model RNA for each step was released by 1 U of nuclease P1 (Wako) at 37 °C and 800 r.p.m in ThermoMixer (eppendorf) for 15 min, then analyzed by HPLC (Agilent 1260 Infinity) using C18 column (5 μm, 150×4.6 mm, Bonna-Agela Technologies).

### Determination of the formation efficiencies of hm^6^A and dm^6^A in poly(A)^+^ RNA using LC-MS/MS analysis

After FTO oxidation and DTT labeling, 200 ng Poly(A)^+^ RNA was digested by 1 U of nuclease P1 (Wako) in 50 μL water solution at 42 °C for 1 h, followed by the addition 1 U of shrimp alkaline phosphatase (rSAP, NEB). The mixture was incubated at 37 °C for an additional 1 h. Digested samples were filtered through 0.22-mm syringe filters prior to UPLC-MS/MS analysis. The nucleosides were separated by UPLC (Shimadzu) equipped with a ZORBAX SB-Aq column (Agilent) and detected by MS/MS using a Triple Quad 5500 (AB SCIEX) mass spectrometer in positive ion mode by multiple reaction monitoring. Nucleosides were quantified using the nucleoside-to-base ion mass transitions of *m/z* 268.0 to 136.0 (A), *m/z* 245.0 to 113.1 (U), *m/z* 244.0 to 112.1 (C), *m/z* 284.0 to 152.0 (G), *m/z* 282.0 to 150.0 (m^6^A), *m/z* 298.1 to 136.0 (hm^6^A), *m/z* 435.0 to 148.1 (dm^6^A). Standard curves were generated by running a concentration series of pure commercial nucleosides and our synthesized nucleoside standards. Concentrations of nucleosides in samples were calculated by fitting the signal intensities to the standard curves.

### Dot-blot assay

1 μg biotin-labeled poly(A)^+^ RNA was spotted on Hybond-N^+^ membrane (Amersham), followed by crosslinking twice under 1500 mJ 254 nm UV (UVP). After dried by air, the membrane was blocked by 1% BSA at room temperature for 1 h and then washed by 1×PBST buffer for three times. HRP-streptavidin (Invitrogen, 1:15000 dilution in PBST) was used to incubate with the membrane at room temperature for 1 h. After washed by PBST for four times, the signals were visualized by Immobilon Western Kit (Millipore) and imaged by Luminescent Imaging Workstation (Tanon 5200).

### m^6^A-SEAL-seq and library construction

5 μg of poly(A)^+^ RNA spiked with 500 pg spike-in RNAs were chemically fragmented into 100-150 nt. The fragmented RNA was treated with FTO oxidation, DTT-mediated thiol-addition reaction, and biotin labeling according to the protocols described above. 50 ng biotin-labeled RNA was saved as input, the rest was proceed to affinity enrichment. 20 μL Dynabeads MyOne Streptavidin C1 (Invitrogen) was washed twice by 200 μL 0.1 M NaOH to remove RNase contamination, then washed with DEPC water to neutral pH. The beads were resuspended in 100 μL binding solution containing 10 μL of high salt wash buffer (100 mM Tris pH 7.5, 10 mM EDTA, 1 M NaCl, 0.05% Tween 20) and 90 μL DEPC water, and incubated with the biotinylated RNA for 1 h. The beads with biotinylated RNA were washed three times with 1 mL high salt wash buffer. 50 μL of 100 mM DTT was used to release the biotinylated RNA at 37 °C for 15 min on ThermoMixer (800 rpm). After collecting the supernatant, the second elution was performed with 50 μL of 100mM DTT at 50 °C for 5 min to completely release the RNA. The twice eluted RNA were combined and purified by by ethanol precipitation. Library construction was performed using NEBNext Ultra II RNA Library Prep Kit for Illumina according to the manufacturer’s protocol. Human libraries were sequenced on Illumina HiSeq XTen platform with pair-end model (PE150). Rice libraries were sequenced on Illumina NextSeq500 platform with sing-end model (SR75).

### FTO-assisted SELECT method

Poly(A)^+^ RNA was treated with FTO following previously published method^30^. FTO-treated and untreated polyA-RNA were mixed with 800fmol Up Primer, 800fmol Down Primer and 1pmol dTTP in 1X CutSmart Buffer (50 mM KAc, 20 mM Tris-HAc, 10 mM MgAc2, 100 μg/ml BSA, pH 7.9 at 25°C). The primers and RNA were annealed by incubating at the following condition: 90°C for 1min; (−1°C/6s)x 41 cycles; 48°C, hold. A 5 μL mixture of 0.001U Bst 2.0 WarmStart DNA Polymerase in 1x CutSmart Buffer was added in the former mixture. The reaction was incubated at 48°C for 20min, and kept at 35°C. Subsequently, a 10 μL mixture containing 0.5U SplintR ligase and 10nmol ATP was added to the final volume 20 μL. The reaction mixture was incubated at 35°C for 15min, denatured at 95°C for 5min, and then kept at 4°C. The quantitative real-time PCR (qPCR) was performed with 2μL reaction mixture as the DNA template. Data was analyzed with QuantStudioTM Real-Time PCR Software v1.3. The detected results of m^6^A site were corrected by neighboring control site.

### Data pre-processing

Reads were trimmed by cutadapt (v1.18) to remove adaptor and reads shorter than 12bp. Trimmed reads were mapped to GRCh38 human genome combining spike-in sequences using HISAT2 (v2.1.0)^42^ Conversion of output sam/bam files were performed by SAMtools^43^ Transcripts from alignment were quantified, assembled and then merged to create a uniform set of transcripts and by StringTie^44^. More precise quantification for each sample was performed by StringTie to obtain FPKM of each gene.

### Identification of m^6^A peaks in transcriptome

Aligned reads were subjected to model-based analysis of ChIP-seq (MACS2) algorithm (v2.1.0)^45^ for peak-calling. The option “--nomodel” and “--slocal 200” was used to call m^6^A peaks and to avoid miss of peaks that shadowed by other adjacent significant peaks. The q-value and enrichment threshold for peaks were 0.05 and 1 respectively. After peak-calling, peaks whose FPKM less than 1 will be filtered to obtain high-confident m^6^A peaks. Overlapped peaks between different samples were found by BEDtools^46^ *intersect* function.

### Motif search

m^6^A peaks were subjected to *De novo* motif search by *findMotifsGenomes.pl* script of HOMER^47^ (v4.10) Output motif length was restricted in 5,6 and 7.

### Metagene profile

Metagene profile was drawn by R package Guitar^48^. Briefly, human annotation GTF file was downloaded from Ensemble (www.ensembl.org). Genomic TxDb object was built from GTF file through *makeTxDbFromGFF* function. Transcriptomic landmarks was built from Genomic TxDb object via *makeGuitarCoordsFromTxDb* function. Peak file was read by *system.file* function and imported as Granges by *import.bed* function. Metagene distribution profile was drawn by transcriptomic landmarks and imported peaks through *GuitarPlot* function.

### Enrichment profile of m^6^A-seal around MeRIP-seq peaks and different motif

Enrichment profile was drawn by deepTools^49^. Briefly, m^6^A-SEAL enrichment score file (bigwig file) was generated from IP and Input bam files through *bamCompare* function. MeRIP-seq region files are bed files from peak-calling, UGUAMM and RRACH region files are generated from rice transcriptome via *scanMotifGenomeWide.pl* script in HOMER package. After that, enrichment signal distribution matrix was calculated by enrichment score files and region files through *ComputeMatrix* function. Enrichment profile and heatmap was generated by enrichment signal distribution matrix through *plotProfile* and *plotHeatmap* function.

### Code availability

Custom Bash, Perl and R codes used for analysis are available on request

