## Supplemental Information for "Antibody-free enzyme-assisted chemical labeling for detection of transcriptome-wide *N*^6^-methyladenosine"

### Supplementary Figures

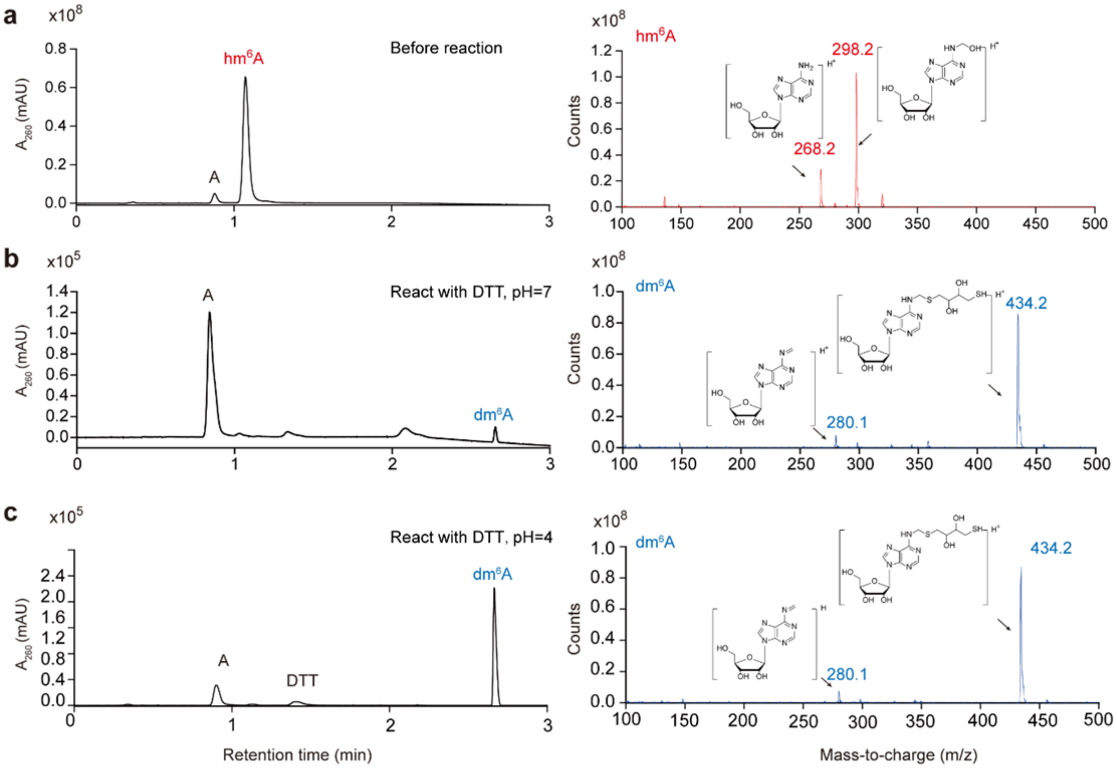


**Supplementary Figure 1. LC-MS characterization of DTT-mediated thiol-addition reaction on hm$^6$A nucleoside.**

Left: HPLC chromatograms. Right: MS profiles of hm$^6$A and dm$^6$A nucleosides. **a**, hm$^6$A before reaction. **b**, The products after hm$^6$A reacted with DTT under pH=7 aqueous solution. **c**, The products after hm$^6$A reacted with DTT under pH=4 aqueous solution.

| Compounds | Yield | Solvent |
|---|---|---|
| 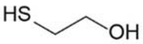  | 95% | $H_2O$ |
| 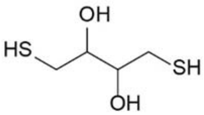  | 72% | $H_2O$ |
| 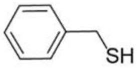  | 42% | MeOH |
| 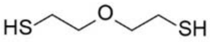  | 32% | MeOH |
| 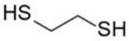  | 26% | MeOH |
| 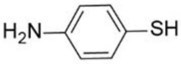  | 25% | MeOH |
| 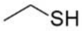  | 16% | MeOH |
| 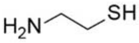  | n.d | $H_2O$ |
| 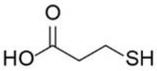  | n.d | $H_2O$ |

**Supplementary Figure 2. Thiol-addition reaction with screening different sulfhydryl-compounds.**

All of the reaction were performed under pH 4, at 37℃ for 3h. The choice of solutions (water or MeOH) are depended on the solubility of individual compounds. Products were characterized by LC-MS.

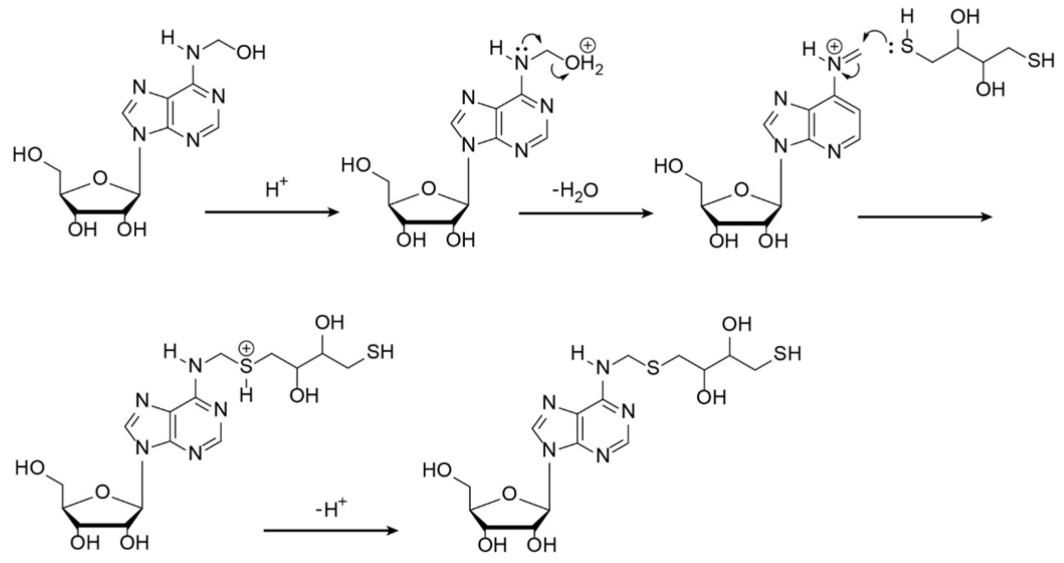


**Supplementary Figure 3. Proposed mechanism for thiol-addition reaction on $hm^6A$.**

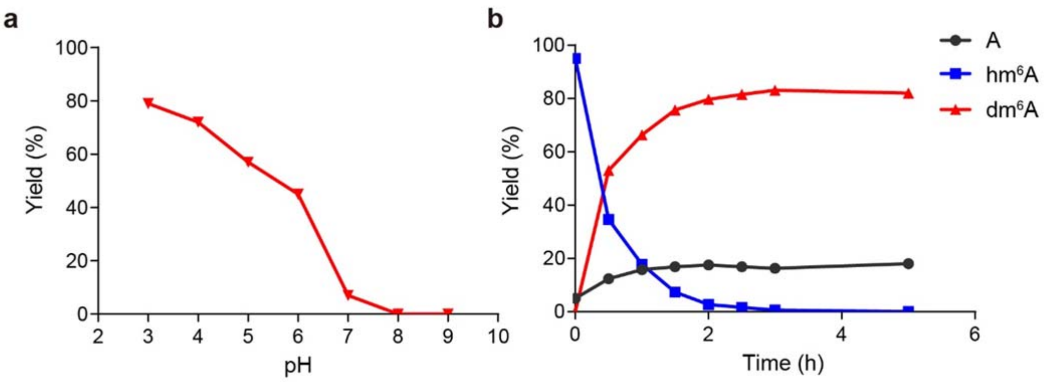


**Supplementary Figure 4. Optimization of DTT-mediated thiol-addition reaction condition on hm$^6$A single nucleoside.**

**a**, Optimization of pH condition for DTT-mediated thiol-addition reaction. 200 mM DTT was used to react with hm$^6$A nucleoside (1 mM) for 3 h in different pH aqueous solution. **b**, Optimization of reaction time for DTT-mediated thiol-addition reaction. 200 mM DTT was used to to react with hm$^6$A nucleoside (1 mM) at pH 4 for different reaction time.

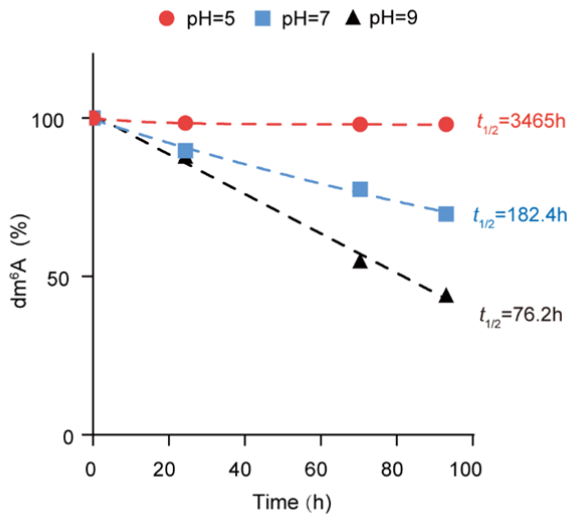


**Supplementary Figure 5. Stability of dm$^6$A nucleoside in aqueous solution with different pH.**

The half-life of dm$^6$A is over 7 days (182 h) in neutral pH, and increases as the pH increases.

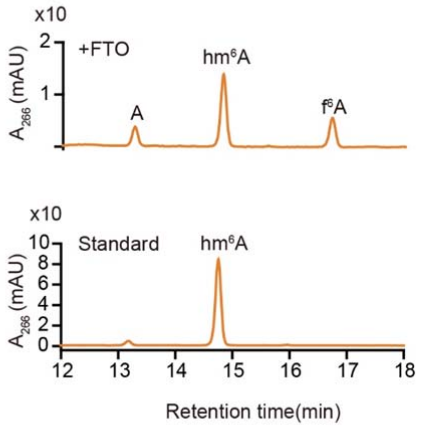


**Supplementary Figure 6. HPLC chromatogram showing FTO oxidation of $m^6A$ to $hm^6A$ in model RNA.**

0.5 nmol of FTO was used to treat with 1 nmol of model RNA for 5 min at pH 7 and 37 °C, generating 60% of $hm^6A$ formation converted from $m^6A$.

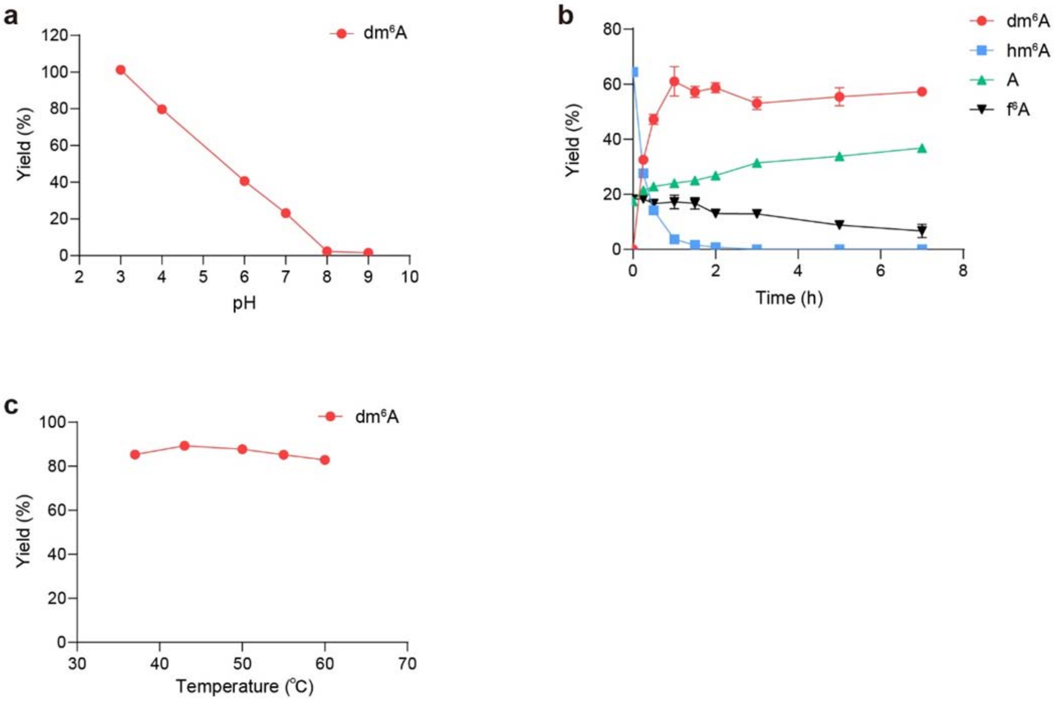


**Supplementary Figure 7. Optimization of DTT-mediated thiol-addition reaction condition on model RNA.**

**a**, Optimization of pH condition for DTT-mediated thiol-addition reaction. 200 mM DTT was used to treat $hm^6A$-modified model RNA for 3 h in different pH aqueous solution. **b**, Optimization of reaction time for DTT-mediated thiol-addition reaction. 200 mM DTT was used to treat $hm^6A$-modified model RNA at pH 4 for different reaction time. **c**, Optimization of reaction temperature for DTT-mediated thiol-addition reaction. 200 mM DTT was used to treat $hm^6A$-modified model RNA at pH 4 for 3 h under different temperature. $hm^6A$-modified model RNA was generated by FTO (0.5 nmol) oxidation of model RNA (1 nmol) 5 min at pH 7 and 37 °C. Data are presented as means ±SD, n=3 biological replicates.

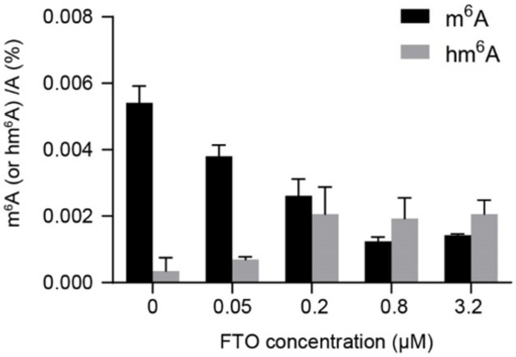


**Supplementary Figure 8. Optimization of FTO usage for the oxidation of $m^6A$ to $hm^6A$ in human poly(A)$^+$ RNA.**

300 ng of poly(A)$^+$ RNA was treated with different concentration of FTO in 300 μL of demethylation reaction for 5 min at 37 °C. Data are presented as means ±SD, n=2 biological replicates × 3 technical replicates.

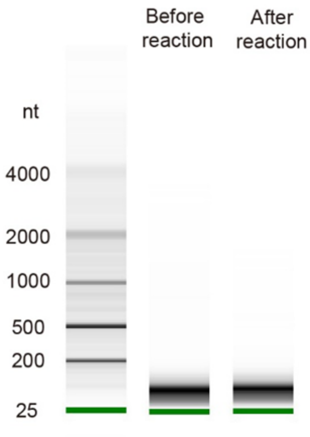


**Supplementary Figure 9. m$^{6}$A-SEAL does not degrade RNA.**

Agilent bioanalyzer result showing no noticeable degradation for the fragmented poly(A)$^{+}$ RNA upon FTO oxidation-assisted DTT thiol-addition reaction.

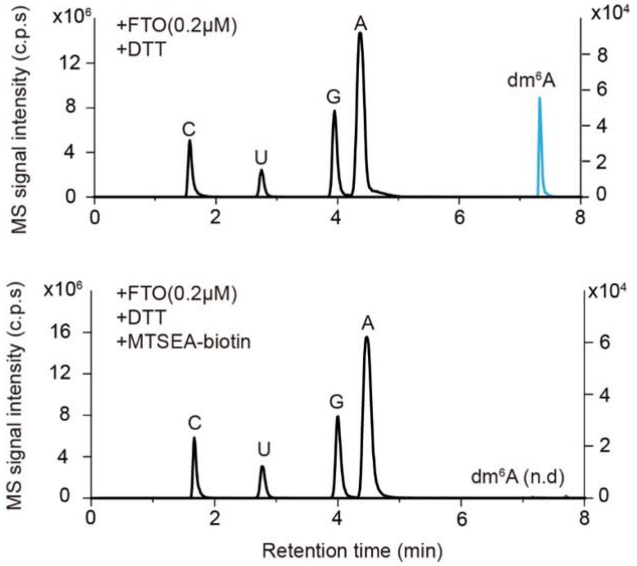


**Supplementary Figure 10. Biotin-labeling efficiency measured by LC-MS/MS.**

LC-MS/MS chromatograms showing no free $dm^6A$ was detected in the RNA sample following the MTSEA-biotin labeling step, indicating very high biotin labeling efficiency.

The chromatograms of A, U, C, and G are scaled to the left y axis, and the chromatograms of $m^6A$, $hm^6A$, and $dm^6A$ are scaled to the right y axis.

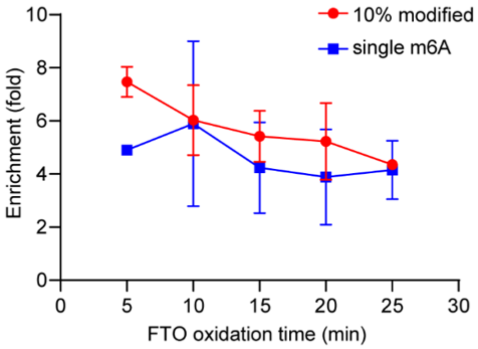


**Supplementary Figure 11. Optimization of FTO oxidation time in $m^6A$-SEAL-seq.**

Enrichment of spike-in RNA sequences under different FTO oxidation time detected by $m^6A$-SEAL-seq. and corresponding enrichment of spike-in RNA. Values represent fold enrichment of IP over input (n=2), normalized to non-$m^6A$-modified spike-in. 10% $m^6A$, reverse transcribed RNA with 10% of $m^6ATP$; single $m^6A$, synthetic RNA with a single $m^6A$ site. 0.2μM FTO was used to oxidize HEK293T poly(A)$^+$ RNA at 37 °C under different reaction time.

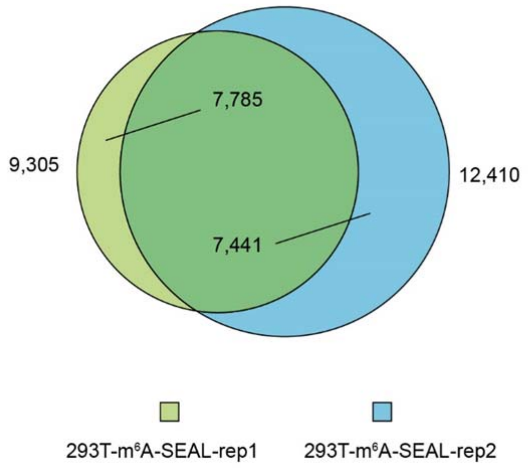


**Supplementary Figure 12. Overlap of m$^6$A sites between two biological replicates for HEK293T cells.**

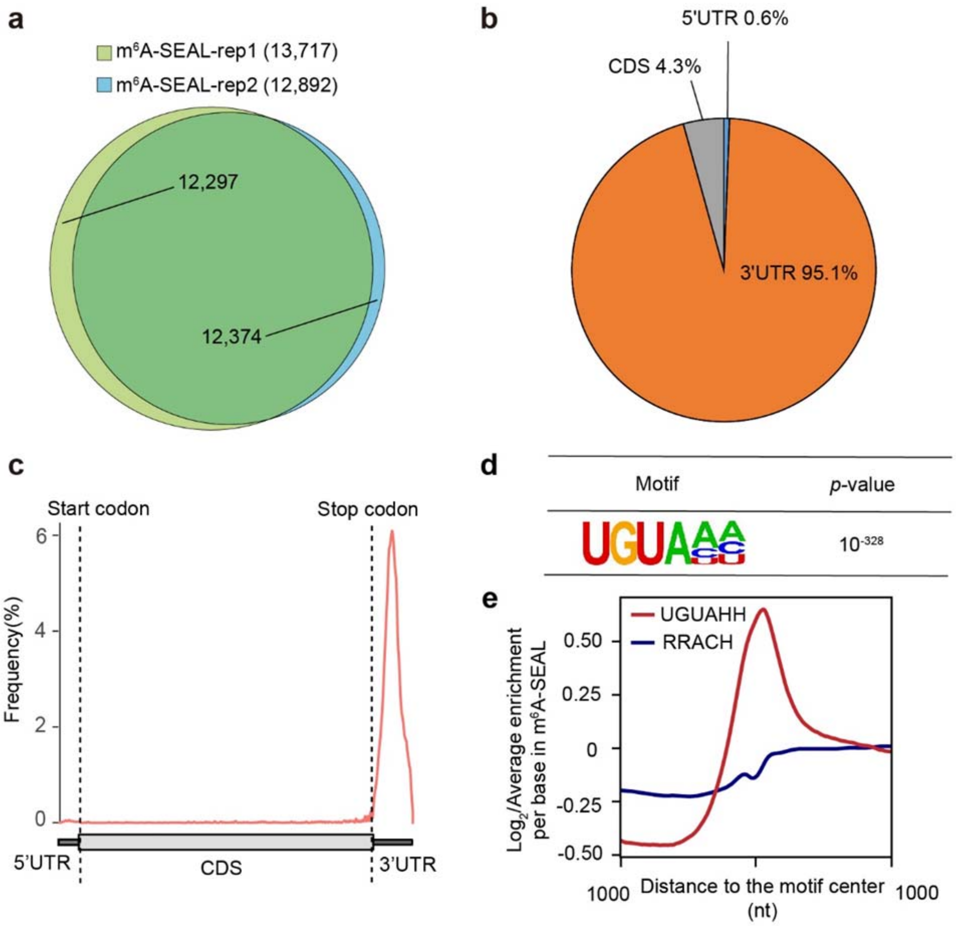


**Supplementary Figure 13. m$^6$A-SEAL-seq uncovers the transcriptome-wide m$^6$A methylome in rice.**

**a**, Overlap of m$^6$A sites between two biological replicates for rice. **b**, Pie chart presenting the fraction of m$^6$A peaks identified by m$^6$A-SEAL-seq in each non-overlapping RNA segment at rice transcriptome. **c**, Metagene profile of m$^6$A sites identified by m$^6$A-SEAL-seq across rice mRNA segments. **d**, A non-canonical rice m$^6$A motif UGUAHH (H=A/C/U) identified in m$^6$A-SEAL-seq. **e**, The enrichment of m$^6$A peaks identified by m$^6$A-SEAL to the centre of UGUAHH or RRACH motifs.

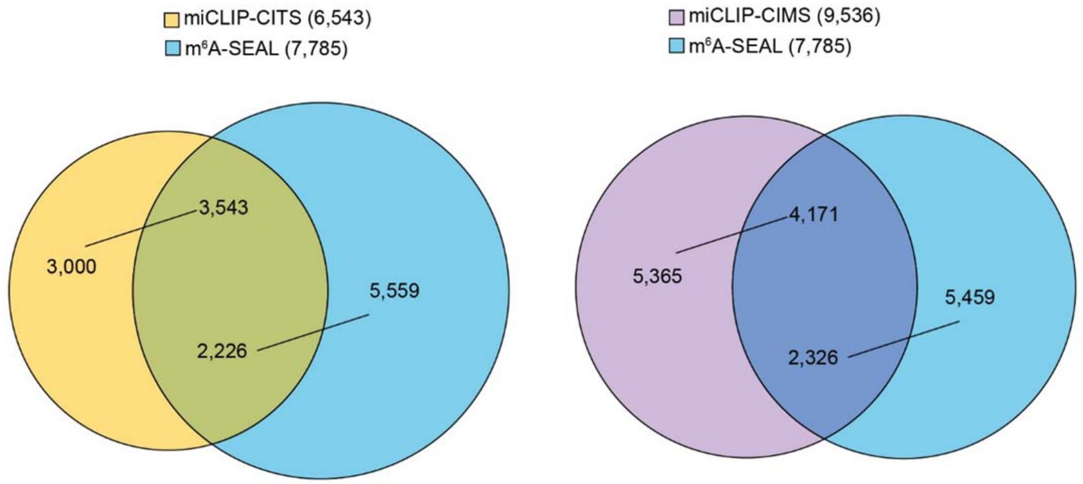


**Supplementary Figure 14. Overlap of identified m$^6$A sites between miCLIP and m$^6$A-SEAL-seq for HEK293T cells.**

Data of miCLIP-CIMS and miCLIP-CITS are obtained from the reference 18.

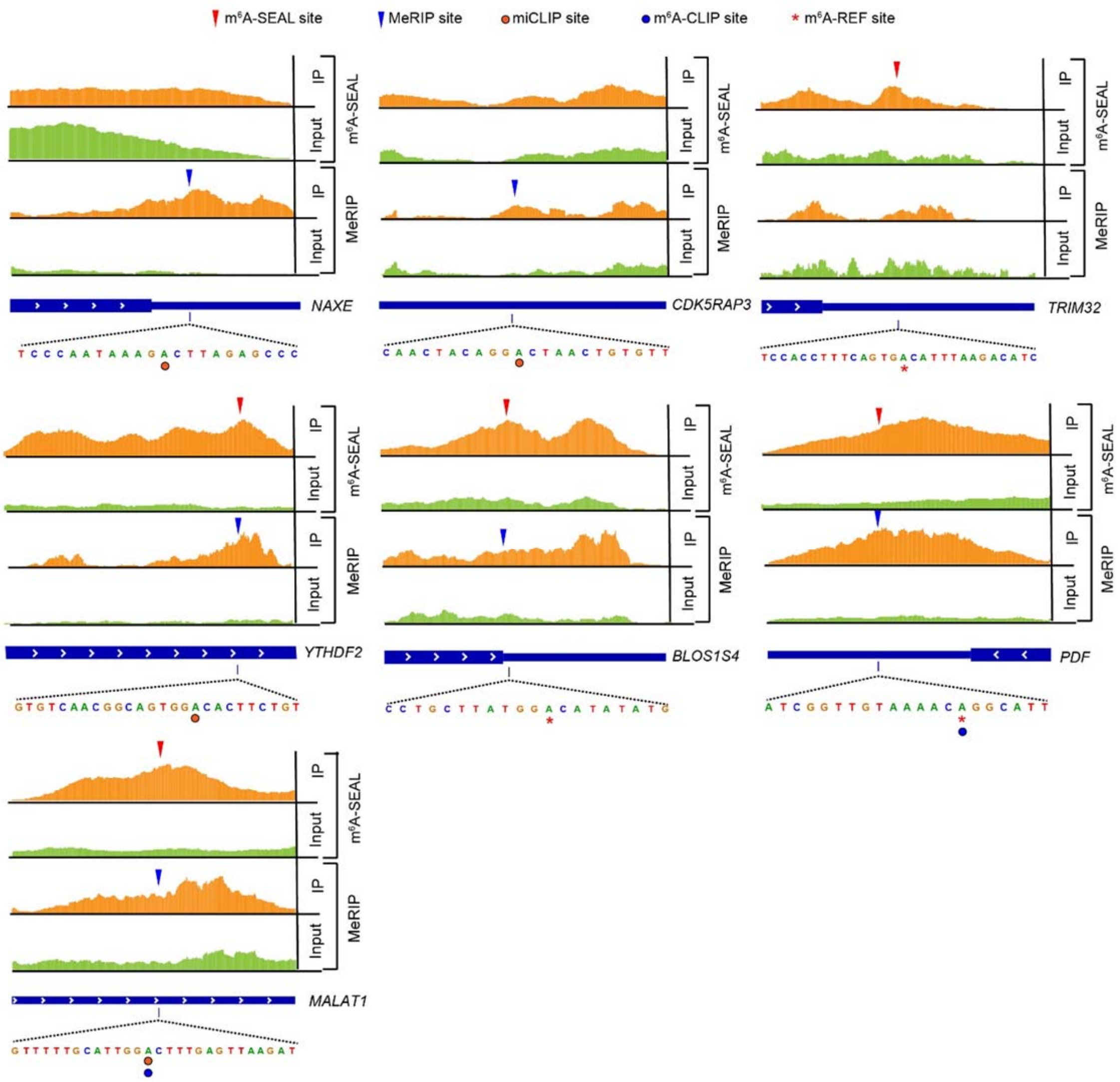


**Supplementary Figure 15. Representative view of m$^6$A sites on mRNAs identified by m$^6$A-SEAL, MeRIP, miCLIP, m$^6$A -CLIP and m$^6$A-REF-seq.**

### Supplementary Notes

**Supplementary Note 1: Preparation and characterization of hm$^6$A single nucleoside**

2 g adenosine nucleoside was suspended in water and treated with 1mL 37% formaldehyde. The mixture was stirred at 60 °C for 10 min and then quenched by 4 mL 20% (w:v) urea solution. hm$^6$A was separated from adenosine by HPLC with XbridgeTM Prep C18 column (Waters).

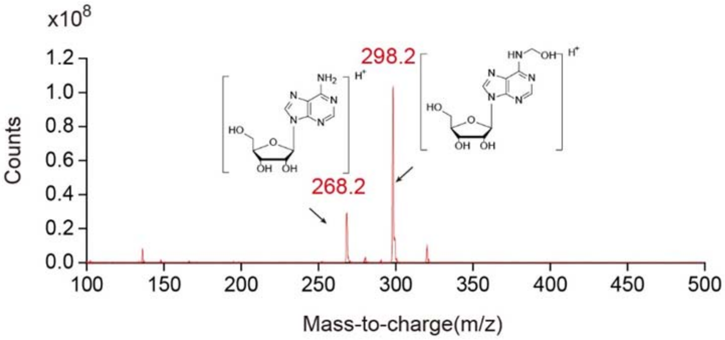


MS characterization of hm$^6$A nucleoside fragment ions:

m/z: 298.2 for [hm$^6$A+H]$^+$, 268.2 for [A+H]$^+$

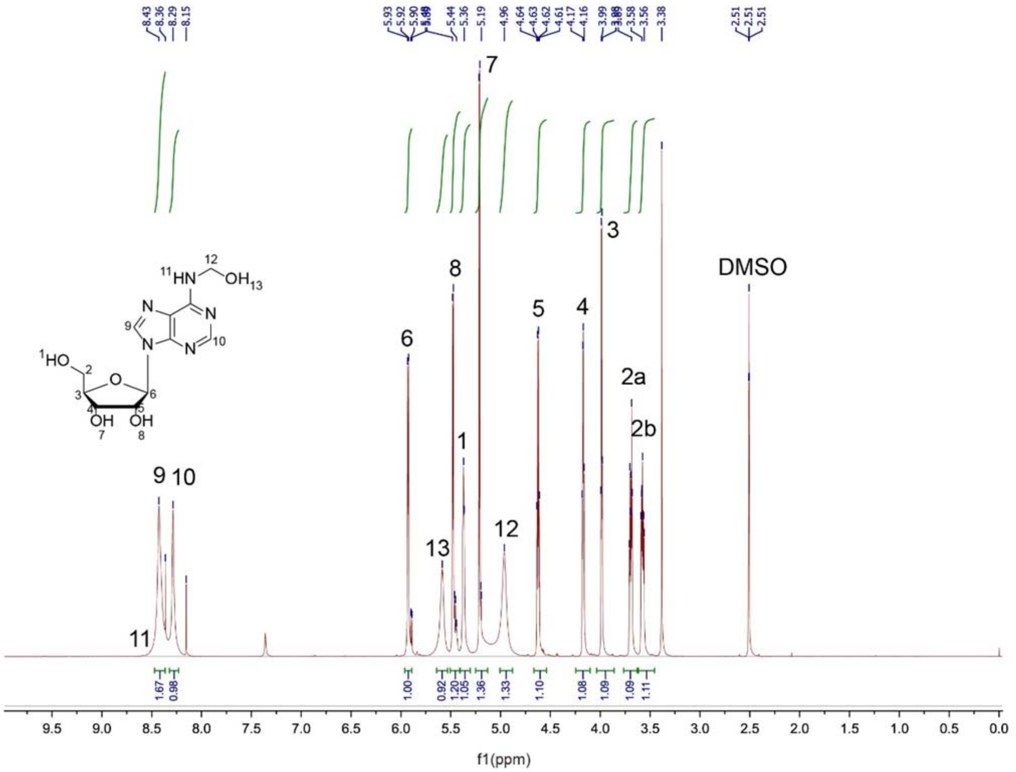


$^{1}$H NMR (700 MHz, DMSO) chart of hm$^{6}$A single nucleoside.

δ ppm: 8.43 (s, 1H), 8.29 (s, 1H), 5.91 (dd, *J* = 23.6, 6.1 Hz, 1H), 5.59 (s, 1H), 5.525.44 (m, 1H), 5.37 (d, *J* = 4.8 Hz, 1H), 5.20 (dd, *J* = 11.4, 4.7 Hz, 1H), 4.96 (s, 1H), 4.62 (dd, *J* = 11.2, 6.0 Hz, 1H), 4.17 (dd, *J* = 8.0, 4.7 Hz, 1H), 3.99 (dd, *J* = 6.8, 3.5 Hz, 1H), 3.69 (dt, *J* = 12.0, 4.1 Hz, 1H), 3.61-3.52 (m, 1H).

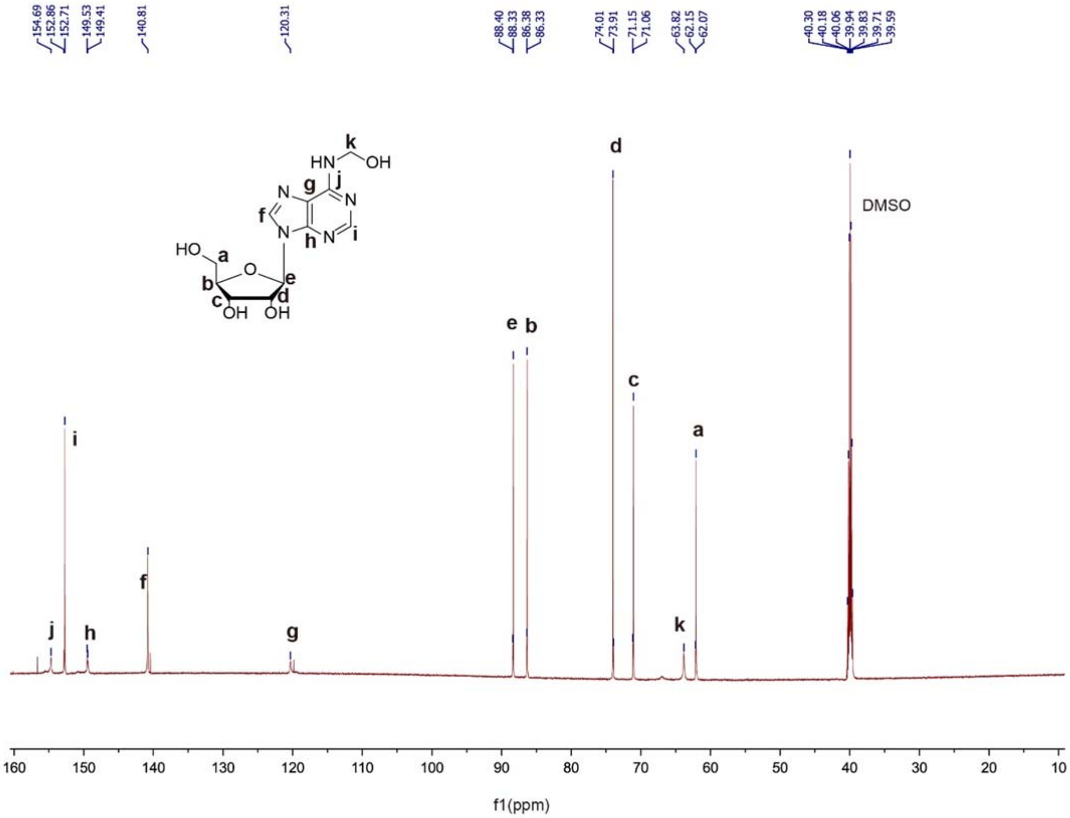


$^{13}C$ NMR (700 MHz, DMSO) chart of $hm^6A$ single nucleoside.

δ ppm: 154.7, 152.7, 149.5, 140.8, 120.2, 88.33, 86.33, 74.0, 71.1, 63.8,62.1,

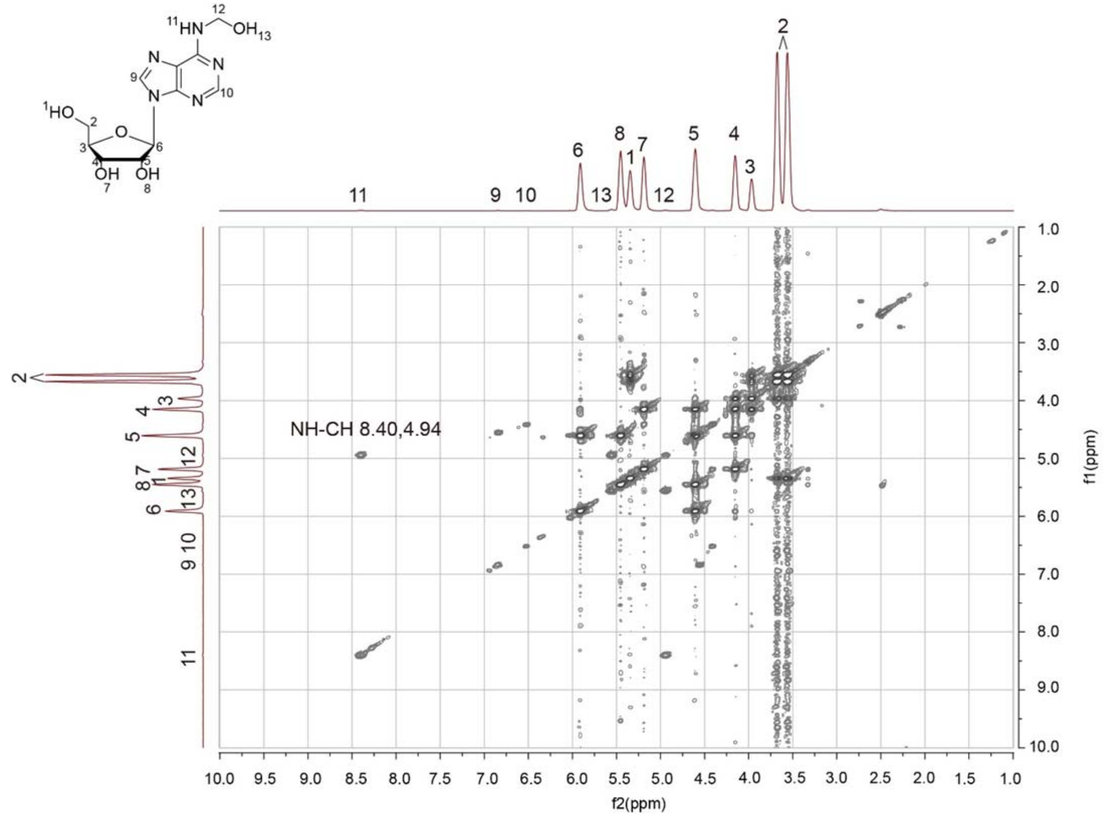


COSY NMR spectrum of hm$^6$A nucleoside.

The correlation between N-H and C-H on 8.40-4.94 for hm$^6$A clearly presented the derivation on exocylic $N^6$ position.

#### Supplementary Note 2: Preparation and characterization of dm$^6$A single nucleoside

500 mg hm$^6$A nucleoside was stirred with 200 mM DTT in 5 mL aqueous solution that contains 100 mM HEPES (pH 4.0) at 37 °C for 3 h, the product dm$^6$A was purified by HPLC with XbridgeTM Prep C18 column and further lyophilized.

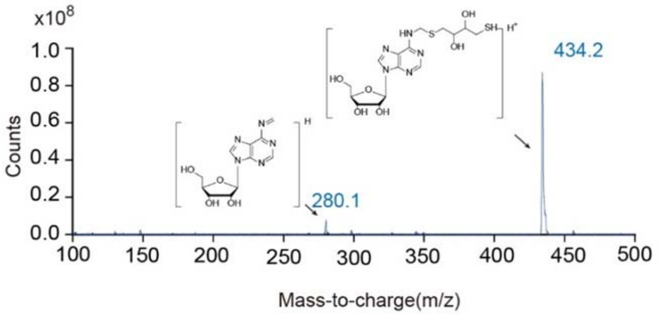


MS/MS characterization of dm$^6$A nucleoside.

Fragment ions m/z: 434.2 for [dm$^6$A+H]$^+$, 280.1 for [*N*$^6$-methyleneadenosine+H]$^+$

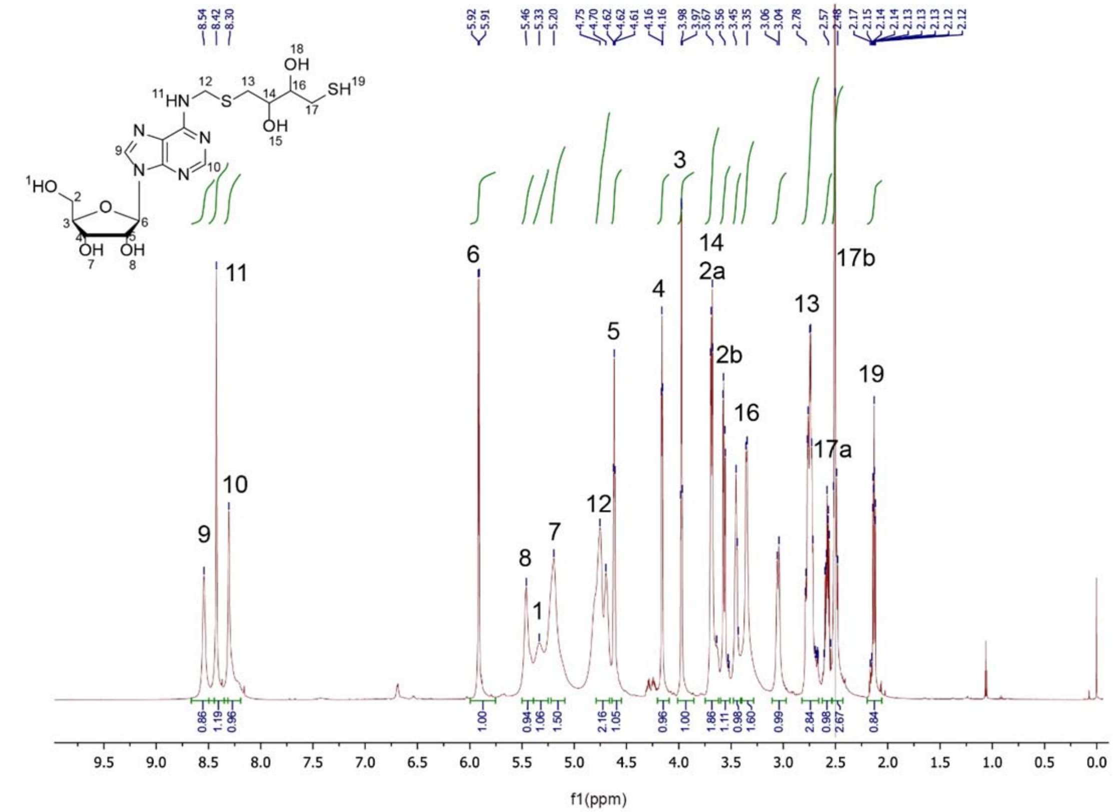


[1]H NMR (700 MHz, DMSO) chart of dm$^6$A single nucleoside.

δ ppm: 8.54 (s, 1H), 8.42 (s, 2H), 8.30 (s, 1H), 5.91 (d, $J$ = 6.0 Hz, 1H), 5.46 (s, 1H), 5.33 (s, 1H), 5.20 (s, 2H), 4.75 (s, 2H), 4.70 (s, 1H), 4.62 (t, $J$ = 5.3 Hz, 2H), 4.22-4.12 (m, 1H), 3.97 (dd, $J$ = 6.8, 3.5 Hz, 1H), 3.68 (dd, $J$ = 12.0, 3.6 Hz, 2H), 3.57 (dd, $J$ = 12.1, 3.6 Hz, 2H), 3.48-3.41 (m, 1H), 3.35 (d, $J$ = 7.9 Hz, 2H), 3.05 (d, $J$ = 12.4 Hz, 1H), 2.87-2.67 (m, 4H), 2.62-2.54 (m, 1H), 2.54-2.40 (m, 4H), 2.18-2.03 (m, 1H).

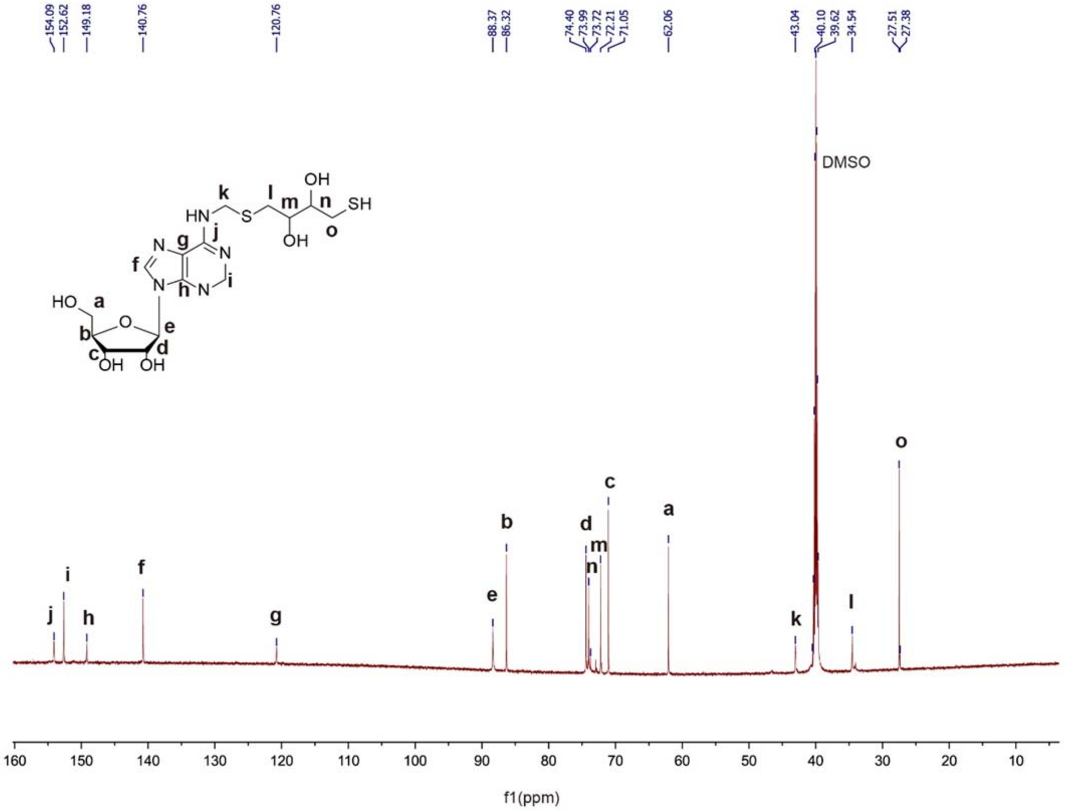


$^{13}$C NMR ( 700 MHz, DMSO) chart of dm$^{6}$A single nucleoside.

δ ppm: 154.1, 152.6, 149.2, 140.8, 120.8, 88.4, 86.3, 74.4, 74.0, 72.2, 71.0, 62.1, 43.0, 34.5, 27.5.

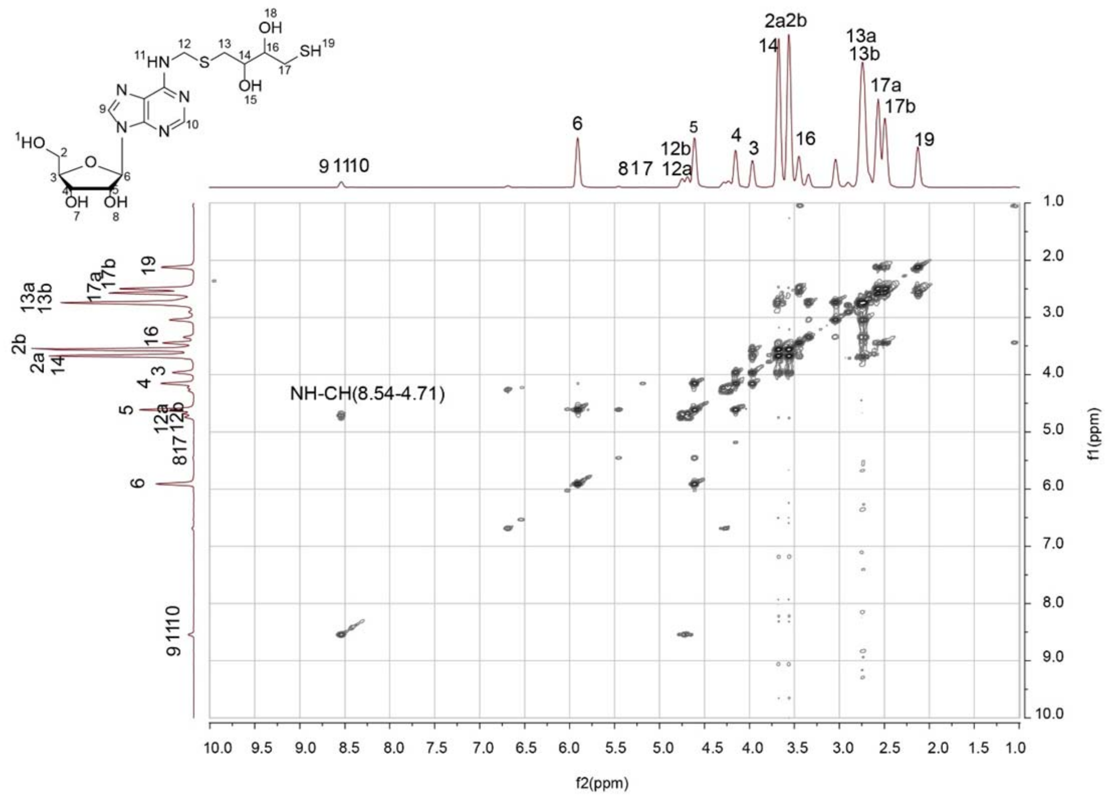


COSY NMR spectrum of dm$^6$A nucleoside.

The correlation between N-H and C-H on 8.54-4.71 for dm$^6$A clearly presented the derivation on exocylic $N^6$ position.

**Supplementary Note 3: Preparation and sequence of spike-in RNAs**

Spike-in RNAs are in-vitro transcribed by HiScribe T7 High Yield RNA Synthesis Kit (NEB) according to the manufacturer's protocol. Briefly, 500ng DNA template was added to the reaction that contains the indicated NTP concentration. 1 µL T7 RNA Polymerase Mix was added to the reaction and the final volume was adjusted to 20 µL. After incubate at 37 °C for 16 h, product RNA was purified by phenol-chloroform extraction. 300 pg modified or unmodified spike in RNAs were added to 3 µg fragmented RNA sample before library preparation.

The DNA template for spike-in RNAs were synthesized by GENERAY. The sequences are as followings. Underlined bases is the promoter of T7 RNA Polymerase and won't be transcribed in reaction.

Unmodified spike-in (7.5 mM CTP, 7.5 mM GTP, 7.5 mM UTP, 7.5 mM ATP)
TAATACGACTCACTATAGGGGTCGGTCGACCTGGGAGCACTGACCCGTATGCTGGA
TCGCCTGGTCTGTAAAGGCTGGGTGGAAAGGTTGCCGAACCCGAATGACAAGCG
CGGCGTACTGGTAAAACTTACCACCGGCGGCGCGGCAATATGTGAACAATGCCAT
CAATTAGTTGGCCAGGACCTGCACCAAGAATTAAC

Single $m^6A$ RNA spike-in (7.5 mM CTP, 7.5 mM GTP, 7.5 mM UTP, 3 mM $m^6ATP$ )
TAATACGACTCACTATAGGGTTTTTGTGTCTTGCCTTTTTCTTTTTTTTGGCTTTT
TGCTTTCCTTCCCTTTCTGTTTGCCGCGTGCCTTCTTTTTCGGGTTTTCCTGACCGC
TGTTCCGTGGGTGTTCTTTCTGTTCTGTGGGGCTTTCGTGGTCGGCTGTCGGGTTC
CCTTTCTTTGGCCCTGTTGGCCGGCCTGCT

10% $m^6A$ spike-in (7.5 mM CTP, 7.5 mM GTP, 7.5 mM UTP, 0.75 mM $m^6ATP$ and 6.75 mM ATP)
TAATACGACTCACTATAGGGATTTTGGACTGGATCGAGGACAACCTGGAATCGCCA
CTGTCACTGGAGAAAGTGTCAGAGCGTTCGGGTTACTCCAAATGGCACCTGCAAC
GGATGTTTAAAAAAGAAACCGGTTTCATCCGCAATAGCAGCTGCATTAATGAATCG
GCCAACGCGCGGGGAGAGGCGGTTTGCGTATTG

### Supplementary Tables

**Supplementary Table 1: Quality and statistics of sequencing**

PE150 model, Hiseq-Xten, Illumina, Human HEK293T cell

| Measure | Replicate1 input | Replicate1 IP | Replicate2 input | Replicate2 IP |
|---|---|---|---|---|
| Total raw reads | 15,351,997 | 16,389,567 | 17,631,193 | 19,170,492 |
| Trimmed reads | 15,351,588 | 16,389,039 | 17,630,189 | 19,169,554 |
| Paired reads | 15,351,588 | 16,389,039 | 17,630,189 | 19,169,554 |
| Mapped paired reads | 13,055,044 | 11,420,876 | 16,416,241 | 16,838,782 |
| Unique mapped paired reads | 11,951,975 | 10,699,217 | 13,065,406 | 14,468,020 |
| Mapping rate in paired reads | 85% | 70% | 93% | 88% |
| Unique mapping rate in paired reads | 78% | 65% | 74% | 75% |

SR75 model, Next-Seq 500, Illumina, Rice Nipponbare

| Measure | Repilcate1 input | Repilcate1 IP | Repilcate2 input | Repilcate2 IP |
|---|---|---|---|---|
| Total raw reads | 24,683,703 | 32,670,382 | 22,846,222 | 29,726,764 |
| Trimmed reads | 24,681,341 | 32,669,573 | 22,843,309 | 29,723,993 |
| Mapped reads | 23,841,001 | 31,518,708 | 21,946,421 | 28,693,542 |
| Uniquely mapped paired reads | 20,283,679 | 28,805,941 | 18,927,741 | 26,413,986 |
| Mapping rate | 96.6% | 96.5% | 96.1% | 96.5% |
| Uniquely mapping rate | 82.2% | 88.2% | 82.8% | 88.9% |

**Supplementary Table 2: List of primers used in SELECT validation of $m^6A$ sites**

| Gene | site | Sequence (5’ to 3’) | Primer |
|---|---|---|---|
| *PNN* | X | TAGCCAGTACCGTAGTGCGTGATCATTGTGCTGATTACCTG | up |
| | | 5phos/CTCCTCCAACTCTTCCTCTCAGAGGCTGAGTCGCTGCAT | down |
| | N | TAGCCAGTACCGTAGTGCGTGTGTGCTGATTACCTGTCTCC | up |
| | | 5phos/CCAACTCTTCCTCTCGCTGAGCAGAGGCTGAGTCGCTGCAT | down |
| *CDK5-RAP3* | X | TAGCCAGTACCGTAGTGCGTGCCCAACCCAAACACAGTTAG | up |
| | | 5phos/CCTGTAGTTGCTCTCAGGCACAGAGGCTGAGTCGCTGCAT | down |
| | N | TAGCCAGTACCGTAGTGCGTGCCCAAACACAGTTAGTCCTG | up |
| | | 5phos/AGTTGCTCTCAGGCACTTGATCAGAGGCTGAGTCGCTGCAT | down |
| *NAXE* | X | TAGCCAGTACCGTAGTGCGTGTGGAAGAGAGGGGCTCTAAG | up |
| | | 5phos/CTTTATTGGGAAGAATACCCACAGAGGCTGAGTCGCTGCAT | down |
| | N | TAGCCAGTACCGTAGTGCGTGGTTCTGGAAGAGAGGGGCTC | up |
| | | 5phos/AAGTCTTTATTGGGAAGAATCAGAGGCTGAGTCGCTGCAT | down |
| *TRIM32* | X | TAGCCAGTACCGTAGTGCGTGACGGGAATATGATGTCTTAAATG | up |
| | | 5phos/CACTGAAAGGTGGAGCAGCAGAGGCTGAGTCGCTGCAT | down |
| | N | TAGCCAGTACCGTAGTGCGTGGGAATATGATGTCTTAAATGTCAC | up |
| | | 5phos/GAAAGGTGGAGCAGATCCCACTCAGAGGCTGAGTCGCTGCAT | down |
| *YTHDF2* | X | TAGCCAGTACCGTAGTGCGTGTCTGCCACGCCACAGAAGTG | up |
| | | 5phos/CCACTGCCGTTGACACTGAACAGAGGCTGAGTCGCTGCAT | down |
| | N | TAGCCAGTACCGTAGTGCGTGACGCCACAGAAGTGTCCAC | up |
| | | 5phos/GCCGTTGACACTGAAAAGTAAGCAGAGGCTGAGTCGCTGCAT | down |
| *BLOC1S4* | X | TAGCCAGTACCGTAGTGCGTGCCACACTCCAACATCATATATG | up |
| | | 5phos/CCATAAGCAGGATTTGCAACAGAGGCTGAGTCGCTGCAT | down |
| | N | TAGCCAGTACCGTAGTGCGTGACTCCAACATCATATATGTCCA | up |
| | | 5phos/AAGCAGGATTTGCAATTCACCAGAGGCTGAGTCGCTGCAT | down |
| *PDF* | X | TAGCCAGTACCGTAGTGCGTGAAGTTGTAGCCAACATTTTG | up |
| | | 5phos/CCGTAACTGATTTCAGGGCACAGAGGCTGAGTCGCTGCAT | down |
| | N | TAGCCAGTACCGTAGTGCGTGTTGTAGCCAACATTTTGTCCG | up |
| | | 5phos/AACTGATTTCAGGGCAAACATTCAGAGGCTGAGTCGCTGCAT | down |
| *MALAT1* | X | TAGCCAGTACCGTAGTGCGTGGGATTTAAAAAATAATCTTAACTCAAAG | up |
| | | 5phos/CCAATGCAAAAACATTAAGTCAGAGGCTGAGTCGCTGCAT | down |
| | N | TAGCCAGTACCGTAGTGCGTGGTCAGCTGTCAATTAATGC | up |
| | | 5phos/AGTCCTCAGGATTTAAAAAATAATCTTAACCAGAGGCTGAGTCGCTGCAT | down |

Note: X site means potential $m^6A$ site; N site means input control site.
